# Blocking Kir6.2 channels with SpTx1 potentiates glucose-stimulated insulin secretion from murine pancreatic β cells and lowers blood glucose in diabetic mice

**DOI:** 10.1101/2022.01.16.476512

**Authors:** Yajamana Ramu, Jayden Yamakaze, Yufeng Zhou, Toshinori Hoshi, Zhe Lu

## Abstract

ATP-sensitive K^+^ (K_ATP_) channels in pancreatic β cells comprise pore-forming subunits (Kir6.2) and modulatory sulfonylurea receptor subunits (SUR1). The ATP sensitivity of these channels enables them to couple metabolic state to insulin secretion in β cells. Antidiabetic sulfonylureas such as glibenclamide target SUR1 and indirectly suppress Kir6.2 activity. Glibenclamide acts as both primary and secondary secretagogues to trigger insulin secretion and potentiate glucose-stimulated insulin secretion, respectively. We tested whether blocking Kir6.2 itself causes the same effects as glibenclamide, and found that the Kir6.2 pore-blocker SpTx1 acts as a strong secondary, but not a primary, secretagogue. SpTx1 triggered a transient rise of plasma insulin and lowered the elevated blood glucose of diabetic mice over-expressing Kir6.2 but did not affect those of non-diabetic mice. This proof-of-concept study suggests that blocking Kir6.2 may serve as an effective treatment for diabetes and other diseases stemming from Kir6.2 hyperactivity that cannot be suppressed with sulfonylureas.

## Introduction

Diabetes mellitus is a group of diseases that all manifest elevated blood glucose levels but with different underlying causes (American Diabetes Association, 2011). Among these diseases, neonatal diabetes mellitus (NDM) was traditionally considered a variant in Type 1 diabetes mellitus (T1DM) and had accordingly been treated with insulin. Since the early 2000s, NDM has been recognized as a genetic disorder that stems from gain-of-function mutations in pancreatic ATP-sensitive K^+^ (K_ATP_) channels (Gloyn et al., 2004). These octameric channel protein complexes, each of which consists of four pore-forming inward-rectifier 6.2 (Kir6.2) subunits and four surrounding auxiliary sulfonylurea-receptor 1 (SUR1) subunits (Aguilar-Bryan et al., 1995; Inagaki et al., 1995), are present in the plasma membrane of β cells within islets of Langerhans in the pancreas. The discovery of the mutations underlying NDM was anticipated by the experimental demonstration in mice that the expression of Kir6.2 with gain-of-function mutations caused hypoinsulinemia and hyperglycemia (Koster et al., 2000). Subsequently, this mutation-caused pathological phenomenon was further demonstrated in mice by over-expressing another gain of function mutant Kir6.2 known to cause NDM at the time (Girard et al., 2009). Mutations in Kir6.2 tend to cause more severe phenotypes of NDM than those in SUR1, and the hyperactivity of the resulting mutant K_ATP_ channels is less likely to be adequately suppressed by sulfonylureas (Hattersley and Ashcroft, 2005; Pipatpolkai et al., 2020). The severe-phenotype-causing mutations of Kir6.2 are also associated with developmental delay, epilepsy and permanent neonatal diabetes (DEND syndrome). In this regard, Kir6.2 represents a new potential drug target.

K_ATP_ channels were originally discovered in the plasma membrane of cardiac myocytes (Noma, 1983) and later found to exist in many other tissue types. K_ATP_ channels in pancreatic β cells are inhibited by both extracellular glucose and intracellular ATP (Ashcroft et al., 1984; Rorsman and Trube, 1985). The ATP sensitivity of these channels is thought to contribute to the regulation of insulin secretion from pancreatic β cells in the following manner (Ashcroft and Rorsman, 2013, 2012; Nichols, 2006). When the blood glucose concentration is low, the overall metabolism in β cells remains low. Consequently, the ratio of intracellular ATP to ADP is relatively low, and the K_ATP_ channels tend to be open, helping to maintain the hyperpolarized resting membrane potential (V_m_). An elevated blood glucose level increases the metabolism in β cells and thus the ratio of intracellular ATP to ADP. A rise of this ratio inhibits K_ATP_ channels, depolarizing V_m_ and thereby increasing the voltage-gated Ca^2+^ channel (Ca_V_) activity. An increased Ca_V_-mediated Ca^2+^ influx raises the concentration of intracellular free Ca^2+^ ([Ca^2+^]_in_), a signal required for triggering robust exocytotic secretion of insulin.

As a class of the antidiabetic drugs, sulfonylureas act as impactful primary secretagogues, triggering insulin release in the presence of a non-stimulating, basal concentration of glucose. These drugs also act as strong second secretagogues, robustly potentiating the insulin secretion stimulated by an elevated concentration of glucose. Sulfonylureas bind to SUR1 and thereby indirectly inhibit currents through the K_ATP_ channels (Gribble and Reimann, 2003; Henquin, 1992). The common sulfonylurea glibenclamide is membrane-permeable and has been shown to lodge inside SUR1, and its effect on insulin secretion cannot be rapidly reversed (Lee et al., 2017; Li et al., 2017; Martin et al., 2017; Schatz et al., 1977; Wu et al., 2018).

The suppression of K_ATP_ activity undoubtedly contributes to the ability of glibenclamide to promote insulin secretion; however, some studies have suggested that glibenclamide also interacts with other proteins involved in the secretory process, in addition to K_ATP_ channels in the plasma membrane (Eliasson et al., 1996; Hinke, 2009; Kang et al., 2011; Lehtihet et al., 2003; Renstrom et al., 2002; Shibasaki et al., 2014; Tian et al., 1998; Zhang et al., 2009). Moreover, while the plasma membrane of β cells is strongly depolarized by raising the concentration of extracellular K^+^ to 30 mM, an application of glibenclamide under this depolarized condition can still promote additional insulin secretion without further changes in [Ca^2+^]_in_ (Geng et al., 2007). These findings about the action of glibenclamide raise the question that to what extent, a direct blockade of the ion-conduction pore of the K_ATP_ channel alone mimics the primary and secondary secretagogue effects of glibenclamide. To address this question, an effective inhibitor of Kir6.2 itself is required.

Our group previously searched for inhibitors of human Kir6.2 (hKir6.2) and discovered that five small proteins in the venoms of certain centipedes inhibited hKir6.2. In particular, a 54-residue protein toxin, dubbed SpTx1, isolated from the venom of *Scolopendra polymorpha*, is the most potent inhibitor with a K_d_ of 15 nM (Ramu et al., 2018; Ramu and Lu, 2019). Here, using SpTx1, we set out to test the impact of direct blocking of Kir6.2 on insulin secretion in mice.

## Results

### SpTx1 does not affect blood glucose levels in wild-type mice

We examined whether SpTx1 had an influence on the resting blood glucose levels of wild-type mice. The blood glucose level of the overnight-fasted wild-type mice was 173 (± 3.18) mg/dL in the following morning (Figure 1A). Given that the blood glucose level of wild-type mice can be lowered by glibenclamide (Remedi and Nichols, 2008), we used it as a positive control. As expected, a single intraperitoneal application of glibenclamide (40 mg/kg body weight) lowered the blood glucose level of wild-type mice by half at the end of a 2-hour observation period whereas that of the vehicle DMSO did not have any meaningful effect. To assess the effects of SpTx1 on blood glucose levels, intravenous (IV) injection was used to administer SpTx1 to avoid any confounding by the toxin bioavailability. SpTx1 at a dose of 1 mg/kg neither had meaningful effect on the blood glucose levels (Figure 1B) of wild-type mice nor caused other noticeable differences during the observation period, compared to wild-type mice administered with vehicle saline. This dose of SpTx1 is calculated to be 3 µM using total blood volume, ∼200 times the K_d_ value of SpTx1 against hKir6.2. The finding that SpTx1, estimated to be at 3 µM in the blood, had no effect on the blood glucose level, suggests that the toxin does not have any consequential actions (in the present context) by either inhibiting mouse Kir6.2 (mKir6.2) channels or acting on unintended targets in wild-type mice.

**Figure 1.**
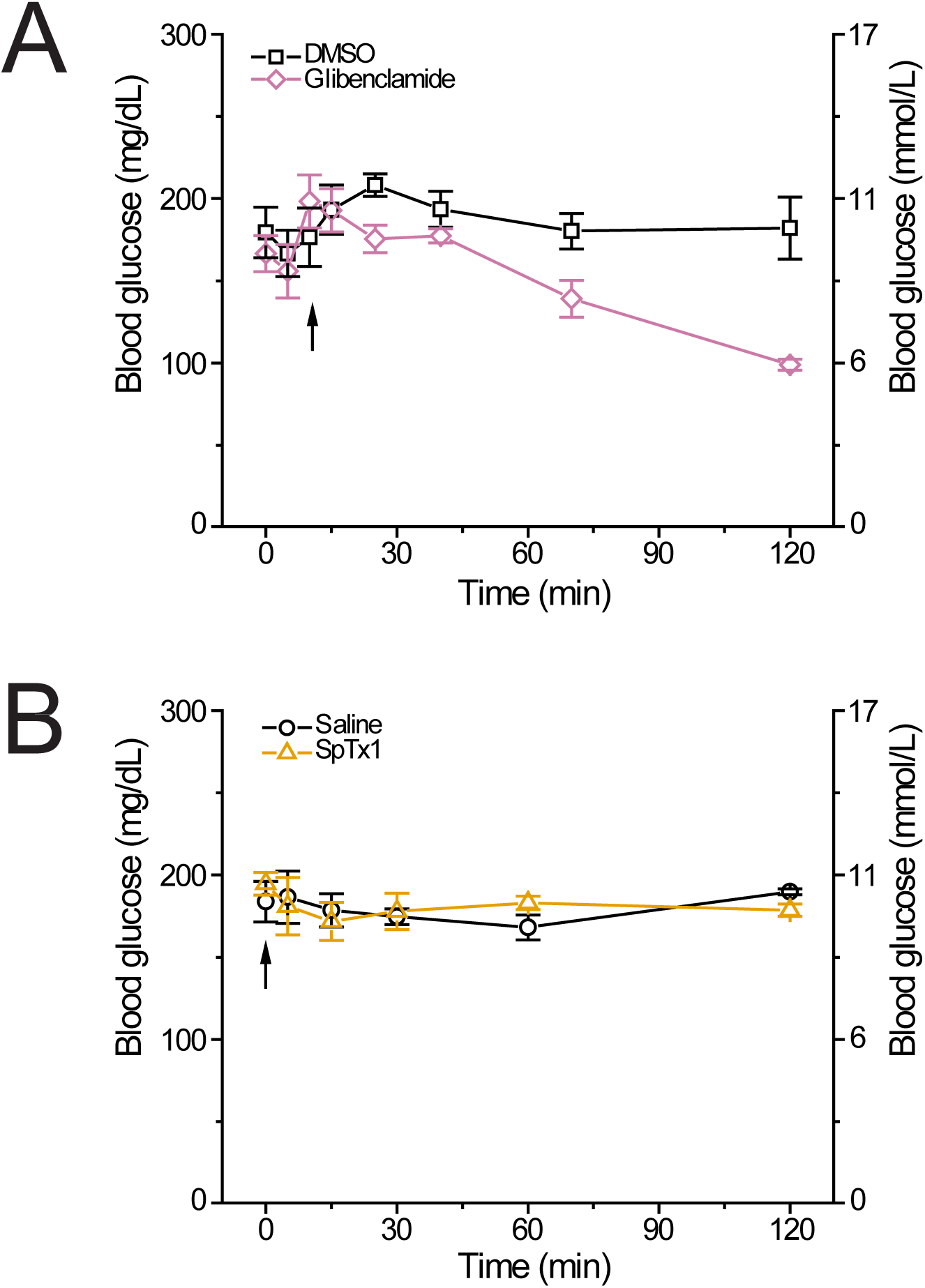
SpTx1 does not affect the blood glucose level of wild-type mice. (**A** and **B**) Blood glucose levels (mean ± SEM, *n* = 5 for each data set) of overnight fasted B6J mice (8 - 12 weeks of age) measured at indicated time points during a 2-hour observation period. (**A**) Glibenclamide (40 mg/kg, purple diamonds) or its vehicle pure DMSO (black squares) was administered by an intraperitoneal injection (arrow). (**B**) SpTx1 (1 mg/kg, orange triangles) or its vehicle saline solution (black circles) was administered by an intravenous injection (arrow).

### SpTx1 fails to potentiate insulin secretion of pancreatic islets from wild-type mice

We tested whether SpTx1 affected the amount of glucose-stimulated insulin secretion (GSIS) from β cells in isolated pancreatic islets of wild-type mice using two different stimulating concentrations of glucose. To perform this test, we adopted a static assay of insulin release from isolated islets incubated in microwell plates (Remedi et al., 2009). We used high concentrations of SpTx1 between 0.2 and 1 μM in the bulk bath solutions because there was likely a SpTx1 concentration gradient from the extracellular solution to the interior of an islet. In this concentration range, SpTx1 failed to alter insulin secretion in the presence of either 7 mM or 15 mM glucose (Figure 2). Thus, SpTx1 had no meaningful effects on GSIS either through acting on mKir6.2 or any unintended targets within the wild-type mouse islets.

**Figure 2.**
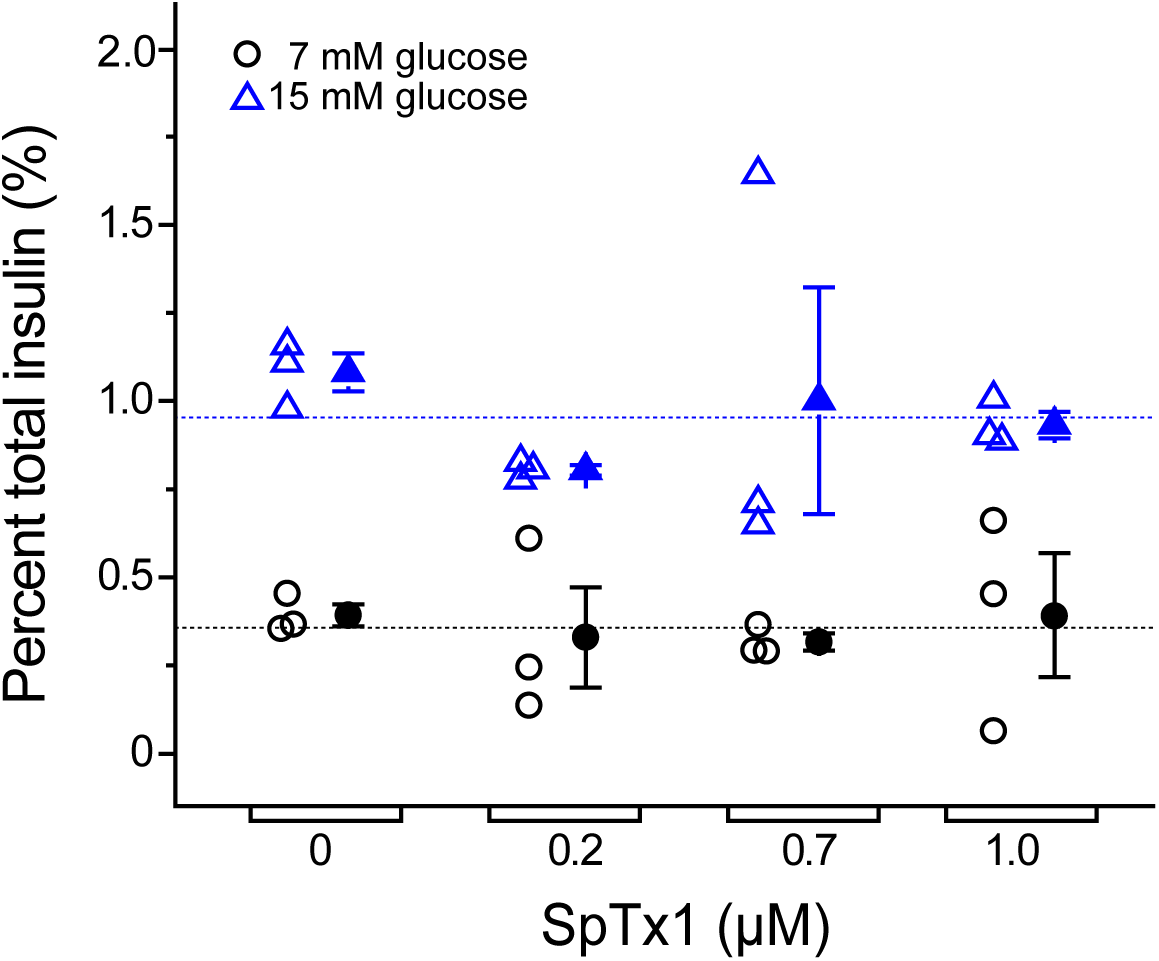
SpTx1 does not potentiate glucose-stimulated insulin secretion from isolated pancreatic islets of wild-type mice. Dot plots of glucose-stimulated insulin secretion from isolated pancreatic islets of B6J mice (8 - 12 weeks of age). For each of the three independent experiments under a given test condition, individual groups of 5 islets incubated in the wells of a microwell plate were used. The secreted insulin as a percentage of the total insulin content of the islets from each assay is plotted against the indicated concentration of SpTx1 in the presence of glucose at the concentration of 7 mM (open black circles) or 15 mM (open blue triangles), with the mean ± SEM of each sample group presented to the right of the respective data set (filled black circles or blue triangles, *n* = 3 independent sets of experiments). The dashed horizonal lines indicate the average of all data in the respective glucose groups.

### SpTx1 does not depolarize the membrane potential of β cells in pancreatic islets from wild-type mice

The prevailing excitation-secretion coupling paradigm of β cell postulates that a blockade of K_ATP_ channels in pancreatic β cells markedly depolarizes V_m_ to trigger insulin release. Previous studies reported that dissociated β cells exhibited altered electrophysiological properties, e.g., a much greater cell-to-cell variation in glucose sensitivity, compared with the cells in isolated islets (Salomon and Meda, 1986; Scarl et al., 2019). Here, V_m_ responses of individual cells in isolated but intact islets were recorded in the perforated whole-cell mode. In a typical mouse islet, up to 80% of the cells are β cells (Steiner et al., 2010). In our measurements, β cells were identified by their characteristic glucose-stimulated V_m_ depolarization and subsequent bursts of (non-overshooting) action potentials. Illustrative V_m_ changes in responses to increasing the extracellular glucose concentration to 15 mM from the starting concentration of 0, 5, or 8 mM are shown in Figure 3A-C, respectively. For example, increasing the glucose concentration from 0 to 15 mM (Figure 3A, left) led to V_m_ depolarization and a burst of action potentials in a reversible manner (Figure 3A, left blue and black segments), thus functionally confirming that the cell recorded from was a β cell. Subsequent application of SpTx1 at 0.2 µM had little effect on V_m_ (Figure 3A, left orange segment) but that of the membrane permeable sulfonylurea glibenclamide at 0.4 µM dramatically depolarized V_m_ and triggered a train of action potentials (Figure 3A, left purple segment). Similar results documenting the ineffectiveness of SpTx1 and the contrasting effectiveness of glibenclamide in inducing V_m_ depolarization were also observed using the starting extracellular glucose concentrations of 5 and 8 mM (Figure 3B and 3C). Low-pass filtered (Figure 3A-C, left light gray traces) time-averaged V_m_ values from multiple islets under the three different conditions are summarized in the right panels of Figure 3A-C. Thus, in wild-type mouse islets, glucose and glibenclamide dramatically depolarize V_m_ but SpTx1, despite its high concentration (>20 times the K_d_ value for hKir6.2), has no such effect. These results also serve as a third control study to determine if SpTx1 has impactful off-targets in the islets.

**Figure 3.**
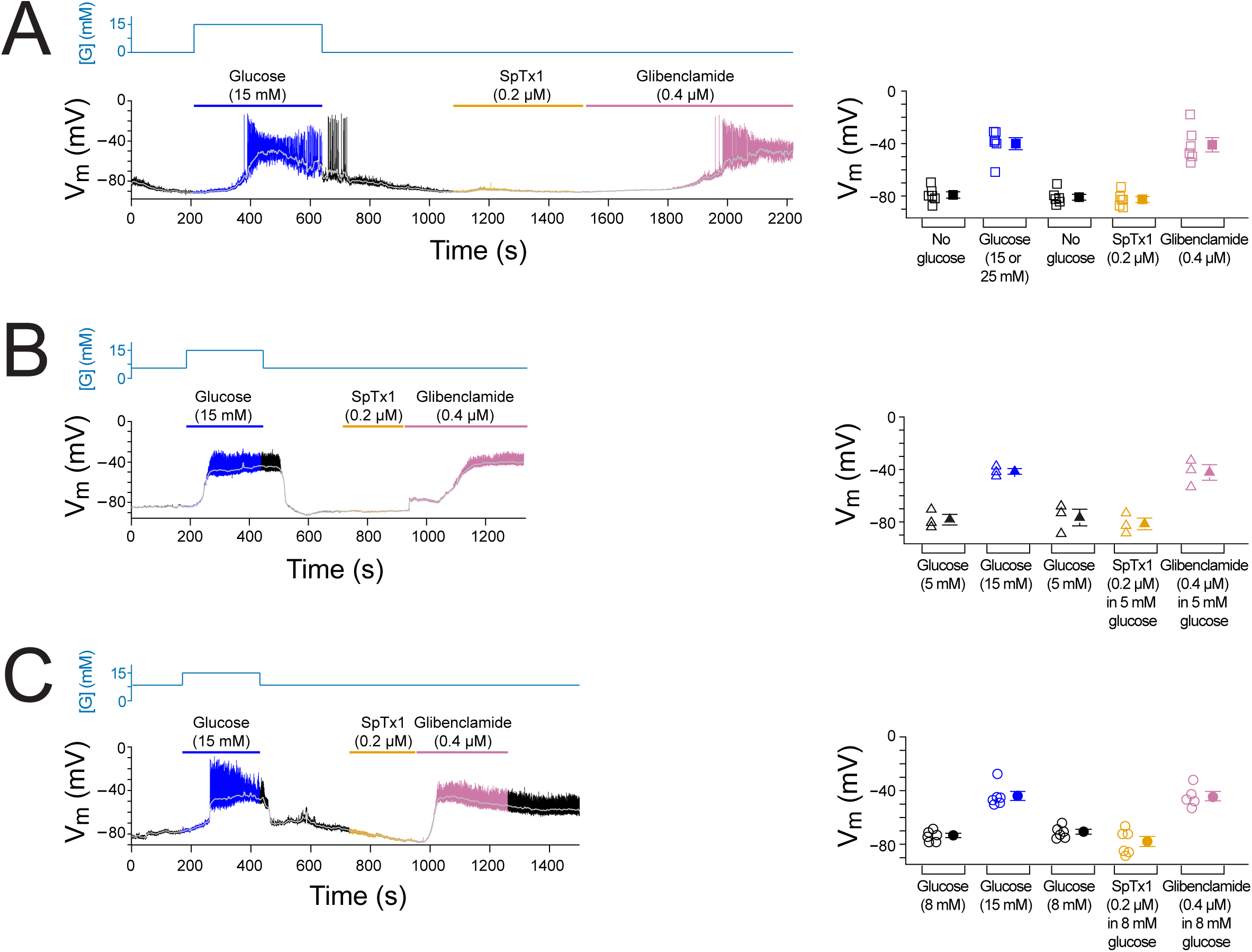
SpTx1 does not depolarize the membrane potential (V_m_) of β cells in isolated pancreatic islets from wild-type mice. (**A**-**C**) Shown on the left are V_m_ traces recorded in the perforated whole-cell mode from individual β cells near the surface of isolated but intact islets from B6J wild-type mice (8 - 12 weeks of age). The switching of the glucose concentration from 0 mM (**A**), 5 mM (**B**) or 8 mM (**C**) to 15 mM (or 25 mM some cases) and back is indicated by the blue schematic line at the top, and the application of 0.2 µM SpTx1 (orange) or 0.4 µM glibenclamide (purple) or the presence of 15 mM (or 25 mM) glucose in the bathing solution (blue) are as indicated by their color-coded lines above the V_m_ trace. The light gray curve overlaid on the V_m_ trace was obtained by (off-line) filtering the recorded trace at 0.1 Hz using a low-pass Gaussian routine. Shown in the right panels are the corresponding resulting values for individual cells under the various conditions, where their mean ± SEM are plotted on the right as filled symbols with errors bars (open triangles, *n* = 6 in **A**; open squares, *n* = 3 in **B**; open circles, *n* = 5 or 6 in **C**).

### SpTx1 inhibits hKir6.2 and mKir6.2 with markedly different affinities

The most likely cause for the failure of SpTx1 to depolarize V_m_ and to potentiate GSIS in wild-type mouse β cells is that SpTx1 does not potently inhibit mKir6.2. Indeed, the amino-acid sequences of hKir6.2 and mKir6.2 are extremely similar but not identical (96% identity). To determine whether SpTx1 targets the two K_ATP_ channel orthologs with different affinities, we examined mKir6.2 and hKir6.2 co-expressed with their respective SUR1 in *Xenopus* oocytes so that we could directly compare the effect of SpTx1 on mK_ATP_ channels with that on hK_ATP_ channels under the same conditions (Figure 4).

**Figure 4.**
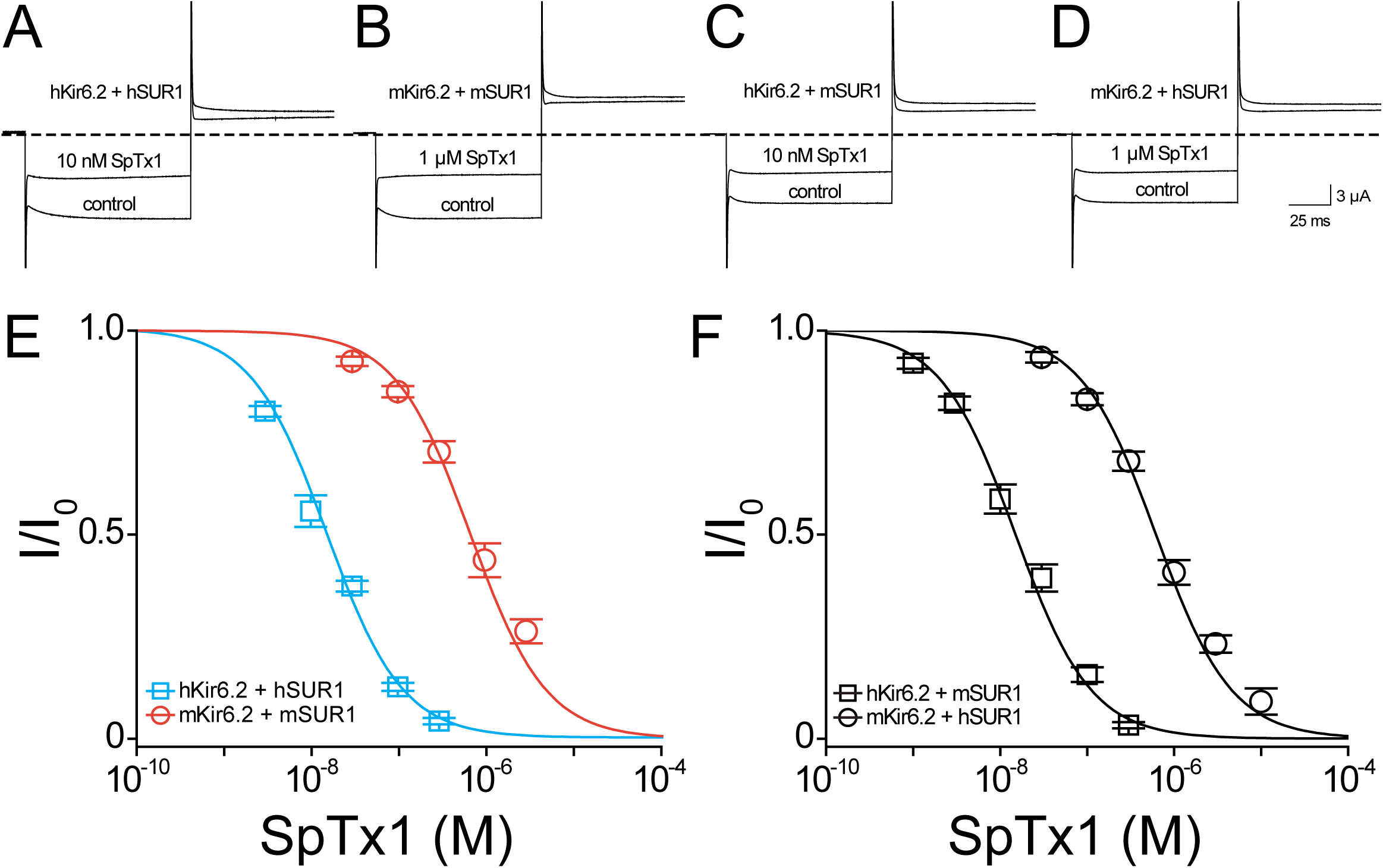
SpTx1 inhibits hKir6.2 and mKir6.2 with markedly different affinities. (**A-D**) Currents of hKir6.2 co-expressed with hSUR1 (hK_ATP_; **A**), mKir6.2 with mSUR1 (mK_ATP_; **B**), hKir6.2 with mSUR1 (**C**), and mKir6.2 with hSUR1 (**D**) activated by adding 3 mM azide to the 100 mM K^+^-containing bath solution and recorded in the absence (control) or presence of 10 nM (**A**, **C**) or 1 μM (**B**, **D**) SpTx1. The currents were elicited by stepping voltages from the holding potential of 0 mV to −80 mV and then to +80 mV. The dashed line indicates zero-current level. (**E** and **F)** Fractions of remaining channel currents (I/I_o_) plotted against the concentration of SpTx1 ([SpTx1]). The curves superimposed on data correspond to the fits of an equation for a bimolecular reaction. The fitted K_d_ values are 1.53 (± 0.13) × 10^-8^ M for hKir6.2 co-expressed with hSUR1 (cyan squares, **E**), 6.27 (± 1.02) × 10^-7^ M for mK_ATP_ with mSUR1 (vermilion circles, **E**), 1.47 (± 0.14) × 10^-8^ M for hKir6.2 with mSUR1 (black squares, **F**), and 6.44 (± 0.69) × 10^-7^ M for mK_ATP_ with hSUR1 (black circles, **F**), where data are plotted as mean ± SEM (*n* = 5 - 8).

As expected, 10 nM SpTx1 suppressed currents through hK_ATP_ channels comprised of hKir6.2 and hSUR1 (hKir6.2 + hSUR1) by about half (Ramu et al., 2018). In contrast, a comparable suppression of current through mK_ATP_ channels, each of which consists of mKir6.2 and mSUR1 (mKir6.2 + mSUR1), required 1 µM SpTx1, a 100 times higher concentration (Figure 4A and 4B). The concentration dependence of current inhibition by SpTx1 was fitted using a model for the toxin-to-channel interaction with one-to-one stoichiometry, yielding an apparent K_d_ of 15 nM and 0.67 μM for hK_ATP_ and mK_ATP_, respectively (Figure 4E). For ease of comparison, all apparent K_d_ values in the entire study are summarized in Table 1. Clearly, SpTx1 inhibits mK_ATP_ with a much lower potency than hK_ATP_.

**Table 1.** SpTx1 affinities for Kir6.2 and its variants.

|  | K<sub>d</sub> in [M] |  |
| --- | --- | --- |
|  | Mean | SEM |
| hKir6.2 + hSUR1 | 1.53173E-8 | 0.12596E-8 |
| mKir6.2 + mSUR1 | 6.72751E-7 | 1.02428E-7 |
| hKir6.2 + mSUR1 | 1.47044E-8 | 0.14003E-8 |
| mKir6.2 + hSUR1 | 6.43655E-7 | 0.69090E-7 |
| hKir6.2<sup>bgd</sup> | 2.07999E-8 | 0.18267E-8 |
| mKir6.2<sup>bgd</sup> | 3.32634E-7 | 0.42909E-7 |
| hKir6.2<sup>bgd</sup>-S103G+A107N+E108V | 3.11112E-7 | 0.33782E-7 |
| hKir6.2<sup>bgd</sup>-S103G+A107N | 2.47986E-8 | 0.28236E-8 |
| hKir6.2<sup>bgd</sup>-S103G+E108V | 3.22783E-7 | 0.38628E-7 |
| hKir6.2<sup>bgd</sup>-A107N+E108V | 3.63467E-7 | 0.43589E-7 |
| hKir6.2<sup>bgd</sup>-S103G | 2.51259E-8 | 0.30150E-8 |
| hKir6.2<sup>bgd</sup>-A107N | 2.93342E-8 | 0.28016E-8 |
| hKir6.2<sup>bgd</sup>-E108V | 2.64264E-7 | 0.33726E-7 |
| mKir6.2<sup>bgd</sup>-V108E | 1.19794E-8 | 0.18805E-8 |
| mKir6.2<sup>bgd</sup>-G103S+N107A+V108E | 1.41949E-8 | 0.25131E-8 |
| hKir6.2-E108V + hSUR1 | 6.15850E-7 | 0.47620E-7 |
| mKir6.2-V108E + mSUR1 | 1.98826E-8 | 0.18643E-8 |
| hKir6.2<sup>bgd</sup> | 1.60624E-8 | 0.13962E-8 |
| hKir6.2<sup>bgd</sup>-N107Y | 2.07997E-8 | 0.18265E-8 |

E denotes exponent of base 10

### Differences between Kir6.2 orthologs underlie their different SpTx1 affinities

It is imperative to exclude the possibility that some differences between the auxiliary hSUR1 and mSUR1 subunits are the primary causes of the aforementioned large difference in apparent SpTx1 affinities of the two orthologous channel complexes. First, we compared the effects of SpTx1 on currents through channels that consisted of hKir6.2 + hSUR1, hKir6.2 + mSUR1, mKir6.2 + mSUR1, and mKir6.2 + hSUR1 (Figure 4A-D). SpTx1 inhibited currents through hKir6.2 + hSUR1 and hKir6.2 + mSUR1 with high affinities but, in contrast, the toxin inhibited currents through mKir6.2 + mSUR1 and mKir6.2 + hSUR1 with low affinities (Figure 4E and 4F). Second, we examined the Kir6.2 mutant that lacks 26 C-terminal residues, dubbed Kir6.2-ΔC26; unlike wild-type Kir6.2, this mutant forms functional channels on the cell surface without needing co-expression with SUR1 (Tucker et al., 1997). For a technical advantage, we used Kir6.2-ΔC26 with an additional mutation at the N-terminal region, V59G, which boosted the apparent expression of the K_ATP_ current (Proks et al., 2004). The resulting constitutively active channels with this double (background) mutation are hereafter referred to as Kir6.2^bgd^, which carry robust currents in whole oocytes that contain millimolar concentrations of ATP. SpTx1 inhibited currents through hKir6.2^bgd^ much more potently than those through mKir6.2^bgd^ (cyan squares versus the orange circles, Figure 5A). The above two lines of results indicate that differences in the two pore-forming orthologous Kir6.2 themselves, but not in their SUR1, primarily underlie the observed differential SpTx1 affinities, and are consistent with the idea that SpTx1 inhibits currents through K_ATP_ channels by interacting with Kir6.2 (Ramu et al., 2018; Ramu and Lu, 2019). Furthermore, the low affinity of SpTx1 for mKir6.2 explains this toxin’s observed ineffectiveness in influencing GSIS from isolated islets (Figure 2) and V_m_ depolarization in individual β cells (Figure 3).

**Figure 5.**
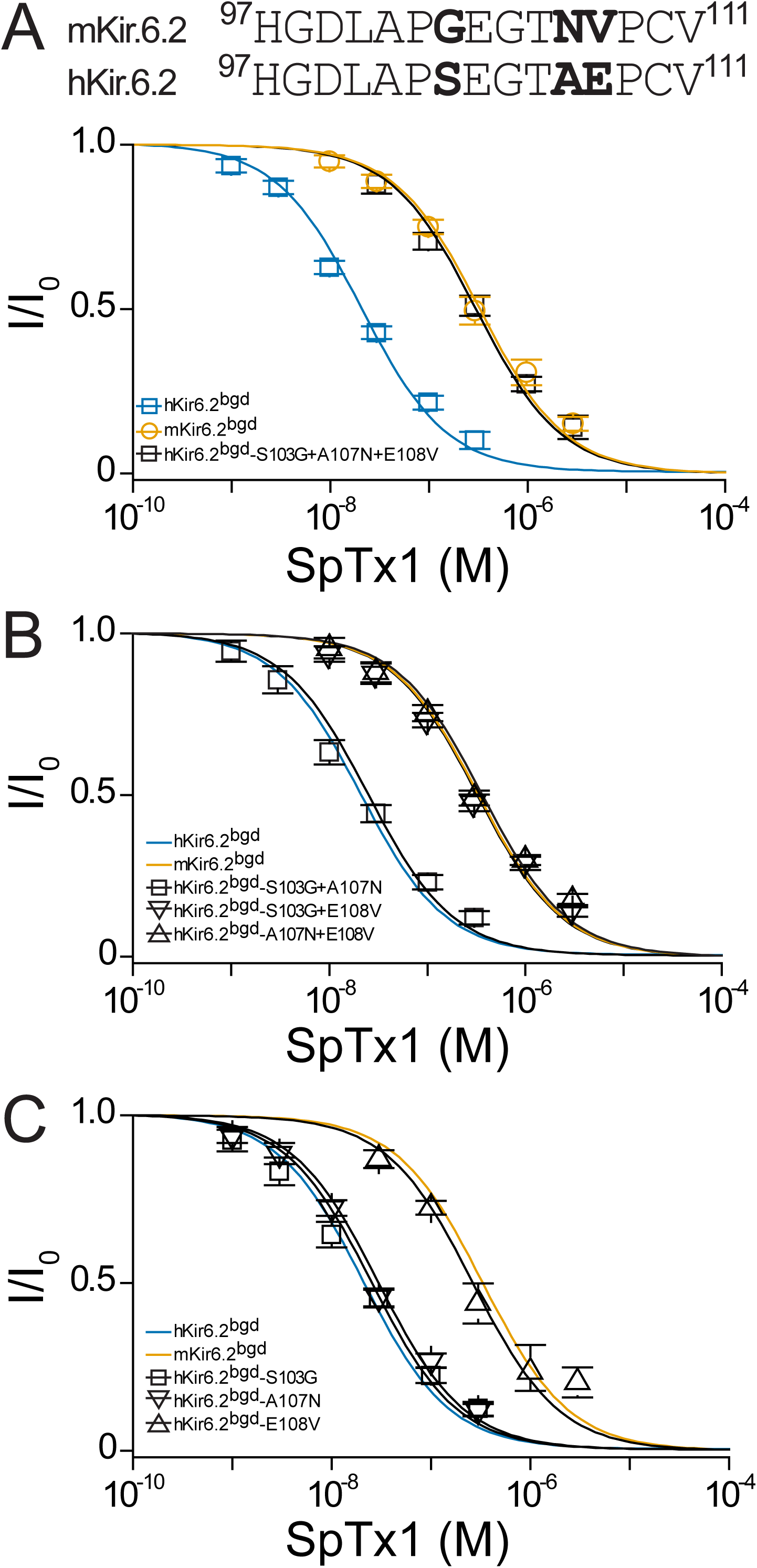
Residue E108 underlies the high affinity of hKir6.2 for SpTx1. (**A**) Shown at top is a comparison of amino acid sequences of a region between the first and second transmembrane segments of hKir6.2 and mKir6.2, which lines the extracellular vestibule of their pore. The three residues that differ between the two sequences are bolded. Shown below are fractions of remaining currents of hKir6.2^bgd^ and mKir6.2^bgd^ plotted against the concentration of SpTx1 along with that of hKir6.2^bgd^ containing an additional triple mutation (S103G+A107N+E108V) to mimic mKir6.2^bgd^. The curves superimposed on data correspond to the fits of an equation for a bimolecular reaction. The fitted K_d_ values are 2.08 (± 0.18) × 10^-8^ M for hKir6.2^bgd^ (blue squares), 3.33 (± 0.43) × 10^-7^ M for mKir6.2^bgd^ (orange circles), and 3.11 (± 0.34) × 10^-7^ M for hKir6.2^bgd^ with the triple mutation (black squares). (**B** and **C**) Fractions of remaining currents of hKir6.2^bgd^ containing additional individual double (**B**) or single (**C**) mutations. The fitted K_d_ values are 2.48 (± 0.28) × 10^-8^ M for hKir6.2^bgd^ containing S103G and A107N (squares), 3.23 (± 0.39) × 10^-7^ M for hKir6.2^bgd^ containing S103G and E108V (inverse triangles), and 3.63 (± 0.44) × 10^-7^ M for hKir6.2^bgd^ containing A107N and E108V (triangles) in **B**, or 2.51 (± 0.30) × 10^-8^ M for hKir6.2^bgd^ containing the mutations of S103G (squares), 2.93 (± 0.28) × 10^-8^ M for hKir6.2^bgd^ containing A107N (inverse triangles), and 2.64 (± 0.34) × 10^-7^ M for hKir6.2^bgd^ containing E108V (triangles) in **C**. For ease of comparison, the fitted curves (blue or orange) for hKir6.2^bgd^ or mKir6.2^bgd^ from **A** are also replotted in **B** and **C**. All data plotted as mean ± SEM (*n* = 5 - 8).

### Variation at a single residue causes different affinities of hKir6.2 and mKir6.2 for SpTx1

Next, we tried to identify the residue variation causing the different SpTx1 sensitivities between hKir6.2 and mKir6.2. Many venom proteins target the external vestibule of the K^+^ channels including Kir channels (MacKinnon et al., 1990; MacKinnon and Miller, 1989; Ramu et al., 2004). Comparison of the sequences of hKir6.2 and mKir6.2 in the region lining the external vestibule of the pore reveals differences at three positions shown in bold (Figure 5A, top). To determine which of the three differences underlies the observed differing SpTx1 affinities, we performed a mutagenesis study with hKir6.2^bgd^ first. The triple mutations (S103G, A107N and E108V), making this segment identical to that in mKir6.2, lowered the affinity of hKir6.2^bgd^ for SpTx1 to the level of mKir6.2^bgd^ (black versus blue squares, Figure 5A).

We then mutated two of the three residues at a time. The SpTx1 affinity of the mutant S103G + A107N (squares, Figure 5B), which did not carry E108V, remained as high as that of hKir6.2^bgd^ itself (blue curve, Figure 5B). In contrast, the remaining two mutants carrying the mutation E108V exhibited markedly reduced SpTx1 affinities (triangles and inverse triangles) similar to that of mKir6.2^bgd^ (orange curve, Figure 5B). These mutagenesis results point to E108V as the mutation responsible for lowering the affinity of hKir6.2^bgd^ for SpTx1. To further confirm this inference, we examined the effects of individual point mutations. Indeed, the E108V mutation alone lowered the affinity of hKir6.2^bgd^ (triangles) to about that of mKir6.2^bgd^ (orange curve) whereas the other two point-mutations had practically no effect on the affinity (squares and inverse triangles), comparable to that of hKir6.2^bgd^ (blue curve, Figure 5C). We also performed the reverse mutagenesis on mKir6.2^bgd^ (Figure 6). Either the triple-mutation (triangles) or the single V108E mutation alone (circles) conferred such a high SpTx1 affinity upon mKir6.2^bgd^ that was comparable to the SpTx1 affinity of hKir6.2^bgd^ (Figure 6; the blue or orange curve is for hKir6.2^bgd^ or mKir6.2^bgd^).

**Figure 6.**
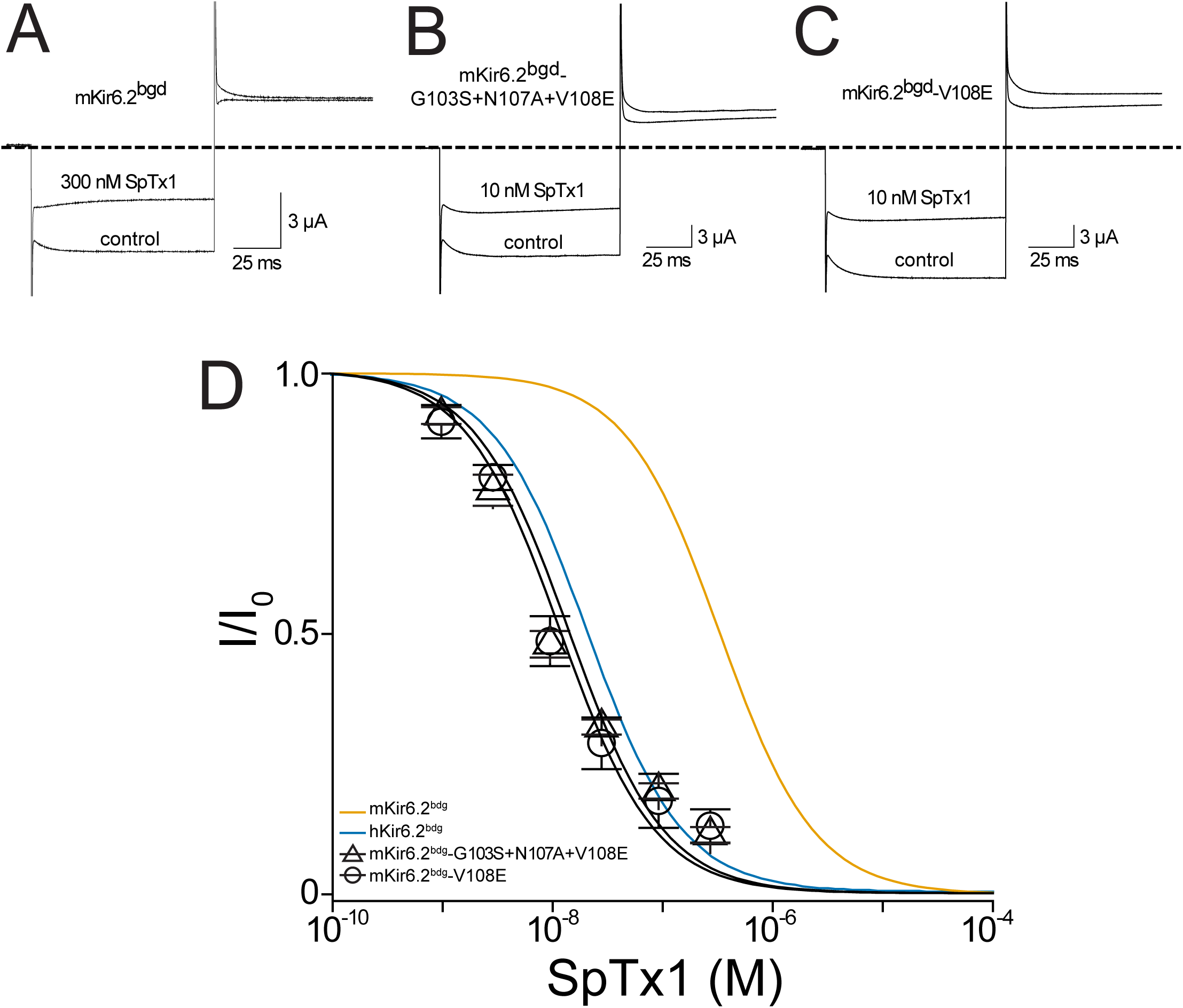
The V108E mutation confers high SpTx1 affinity upon mKir6.2^bgd^. (**A**-**C**) Currents of mKir6.2^bgd^ (**A**), mKir6.2^bgd^ containing the triple mutations G103S, N107A and V108E (**B**), or mKir6.2^bgd^ containing the single mutation V108E (**C**), elicited by stepping voltages from the holding potential of 0 mV to −80 mV and then to +80 mV in the presence of 100 mM K^+^ in the bath solution, and recorded in the absence (control) or presence of 300 nM (**A**) or 10 nM (**B**, **C**) SpTx1. The dashed line indicates zero-current level. (**D**) Fractions of remaining channel currents plotted against the concentration of SpTx1. The curves superimposed on data correspond to the fits of an equation for a bimolecular reaction. The fitted K_d_ values are 1.42 (± 0.25) × 10^-8^ M for mKir6.2^bgd^ containing the triple mutations (triangles) and 1.20 (± 0.19) × 10^-8^ M for mKir6.2^bgd^ containing V108E (circles), where data are plotted as mean ± SEM (*n* = 5 - 8). For ease of comparison, the fitted curves (blue or orange) for mKir6.2^bgd^ or mKir6.2^bgd^ are also replotted from Figure 5A.

To demonstrate that the above results also occur in the octameric K_ATP_ channel-protein complex, we examined hKir6.2 with the E108V mutation and mKir6.2 with the V108E mutation, co-expressed with their respective SUR1. Indeed, the mKir6.2-V108E mutation conferred the high SpTx1 affinity of hK_ATP_ channels upon mK_ATP_ channels co-expressed with mSUR1 (circles, Figure 7). Conversely, the hKir6.2-E108V mutation conferred the low SpTx1 affinity of mK_ATP_ channels upon hK_ATP_ channels co-expressed with hSUR1 (squares, Figure 7).

**Figure 7.**
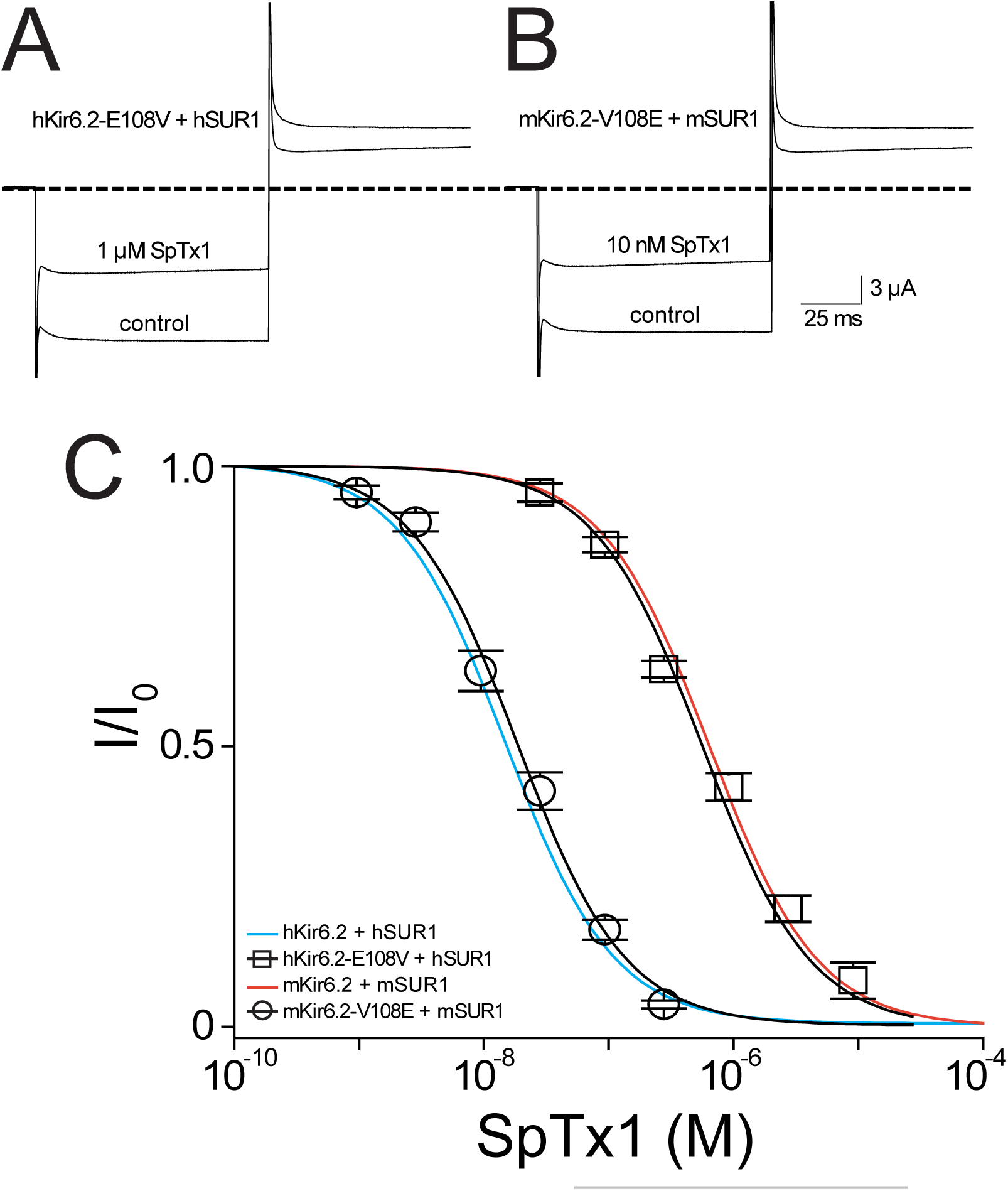
Single point mutations switch the affinities of hK_ATP_ and mK_ATP_ channels for SpTx1. (**A** and **B**) Currents of hKir6.2-E108V co-expressed with hSUR1 (A) or mKir6.2-V108E with mSUR1 (**B**), elicited by stepping voltages from the holding potential of 0 mV to −80 mV and then to +80 mV in the presence of 100 mM K^+^ in the bath, and recorded in the absence (control) or presence of 1 μM (**A**) or 10 nM (**B**) SpTx1. The dashed line indicates zero-current level. (**C**) Fractions of remaining currents of the two mutant channels plotted against the concentration of SpTx1. The fitted K_d_ values are 6.16 (± 0.48) × 10^-7^ M for hK_ATP_-E108V co-expressed with hSUR1 (squares), and 1.99 (± 0.19) × 10^-8^ M for mK_ATP_-V108E with mSUR1 (circles). For ease of comparison, the fitted curves (cyan and vermilion) for hKir6.2 with hSUR1 and mKir6.2 with mSUR1 are also replotted from Figure 4E. All data are plotted as mean ± SEM (*n* = 5 - 8).

The above results demonstrate that the low affinity of mK_ATP_ channels for SpTx1 reflects a residue variant, valine versus glutamate, at amino-acid position 108 located in the extracellular vestibule of the pore. This finding in turn strengthens the notion that SpTx1 inhibits the K_ATP_ channel by blocking Kir6.2’s ion-conduction pore (Ramu et al., 2018; Ramu and Lu, 2019).

### SpTx1 depolarizes the membrane potential of β cells in pancreatic islets from V108E-mutant mice

On the basis of the mechanistic information revealed by the above heterologous mutagenesis studies, we generated a knock-in mutant mouse line, in which the valine 108 residue of the endogenous (Endo) mKir6.2 was replaced by a glutamate residue, dubbed ^Endo^mKir6.2^V108E^. According to the prevailing paradigm, if the high SpTx1 sensitivity is conferred upon the mK_ATP_ channels of ^Endo^mKir6.2^V108E^ mice, then SpTx1 should depolarize V_m_ of their β cells and possibly elicit action potentials.

As performed with wild-type islet cells (Figure 3), we recorded V_m_ from individual β cells in isolated but intact ^Endo^mKir6.2^V108E^ islets (Figure 8). Starting from 0, 5, and 8 mM glucose in the extracellular medium, the islets were challenged with high glucose, SpTx1, or glibenclamide. In a nominal glucose-free condition, SpTx1 at 0.2 µM, 10 times higher than the K_d_ value for V108E-carrying mKir6.2 (+ mSUR1), caused a small but noticeable 7 mV depolarization (orange, Figure 8A). However, this SpTx1-induced V_m_ depolarization was much smaller than that induced by high glucose or glibenclamide that triggered action potentials (blue or pink, Figure 8A). With 5 mM glucose (orange, Figure 8B), SpTx1 at 0.2 µM also caused V_m_ depolarization without eliciting any action potentials (orange, Figure 8B), which was again less than that induced by high glucose or glibenclamide (blue or pink, Figure 8B). However, with 8 mM glucose, SpTx1 at 0.2 µM more strongly depolarized V_m_ and elicited action potentials (orange, Figure 8C). The results pooled from the measurements under various conditions are summarized in the respective panels on the right in Figure 8. Thus, the impact of SpTx1 on V_m_ depends on the glucose concentration. [We note that under the 5 mM glucose condition, SpTx1 had variable effect on V_m_ of β cells in multiple islets (right panel, Figure 8) but the toxin did not elicit any action potentials regardless of the magnitude of apparent depolarization.]

**Figure 8.**
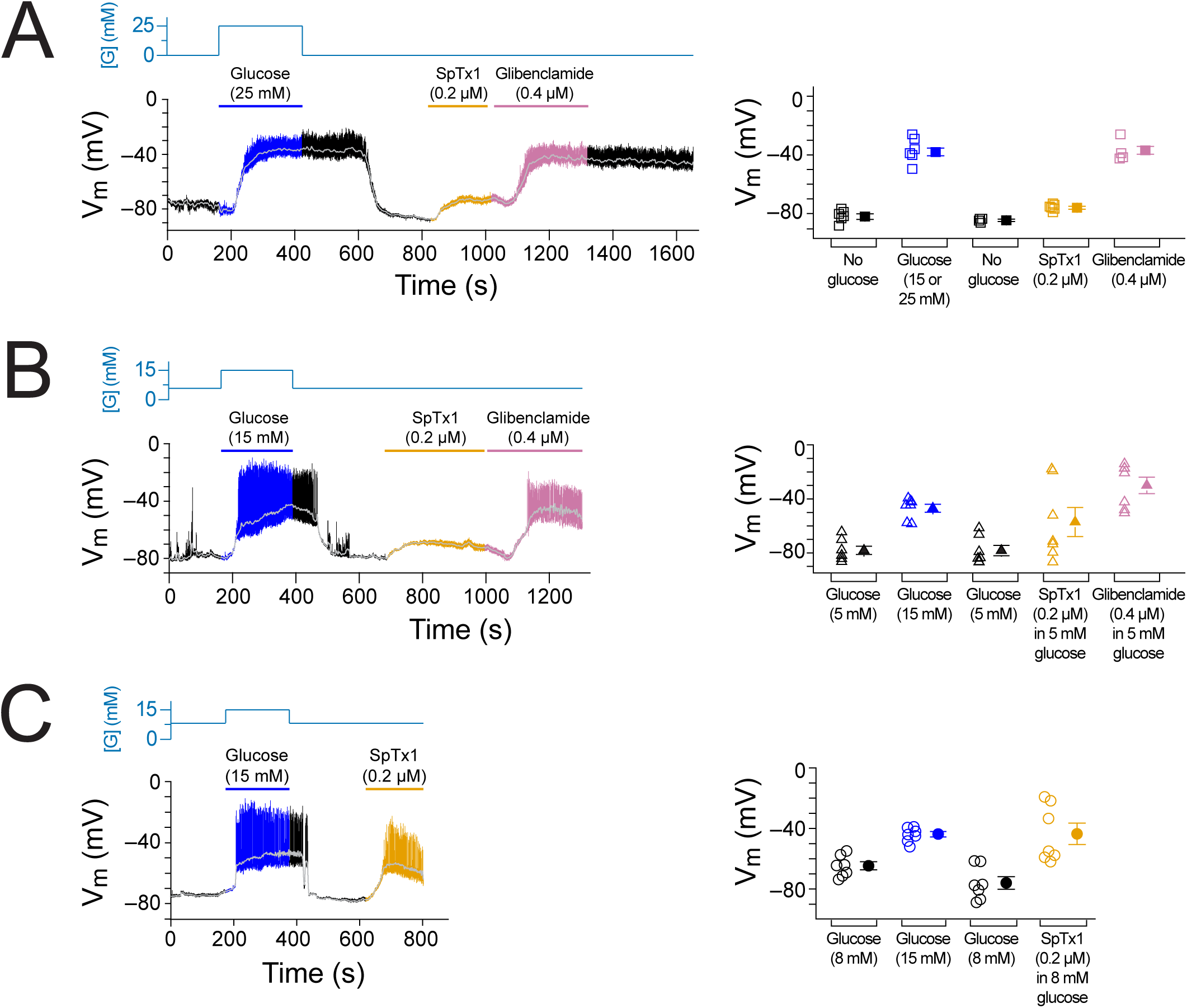
SpTx1 depolarizes the membrane potential (V_m_) of β cells in isolated pancreatic islets from ^Endo^mKir6.2^V108E^ mice. (**A**-**C**) Shown on the right are V_m_ traces recorded in the perforated whole-cell mode from individual β cells near the surface of isolated intact islets from ^Endo^mKir6.2^V108E^ mice (8 - 12 weeks of age). The switching of the glucose concentration from 0 mM (**A**), 5 mM (**B**) or 8 mM (**C**) to 15 or 25 mM and back is indicated by the blue schematic line at the top, and the application of 0.2 µM SpTx1 (orange) or 0.4 µM glibenclamide (purple) or the presence of 15 or 25 mM glucose in the bathing solution are as indicated by their color-coded lines above the V_m_ trace. The light gray curve overlaid on the V_m_ trace was obtained by filtering the recorded trace at 0.1 Hz using a low-pass Gaussian routine. Shown in the right panels are the resulting values for individual cells under the various conditions, where their mean ± SEM are plotted on the right as filled symbols with errors bars (open squares, *n* = 4 or 6) in **A**; open triangles, *n* = 6 in **B**; open circles, *n* = 7 in **C**).

### SpTx1 markedly potentiates GSIS from pancreatic islets of the V108E-mutant mice

We examined GSIS of islets isolated from the pancreas of the ^Endo^mKir6.2^V108E^ mice by a perifusion method. Given that SpTx1 in 0 mM and 5 mM glucose did not trigger action potentials but it did so in 8 mM glucose, we set the basal glucose level to 3 mM, a known non-stimulating concentration for isolated intact islets, and then raised the glucose concentration to 10 mM to stimulate insulin secretion in the initial set of studies. As a positive control, upon elevating the glucose concentration in the perifusion solution from 3 mM to 10 mM, insulin secretion rapidly increased in the first phase and then declined in the second phase to a relatively steady level above the baseline level (black trace, Figure 9A). After the glucose concentration was lowered back to 3 mM, insulin secretion returned to the baseline level. As a further control, elevation of glucose and inclusion of 1 μM glibenclamide together yielded insulin secretion much greater than that from elevated glucose alone. As expected, even after removing glibenclamide from the perifusion solution and lowering glucose back to 3 mM, the elevated insulin secretion largely persisted (cyan trace, Figure 9A).

**Figure 9.**
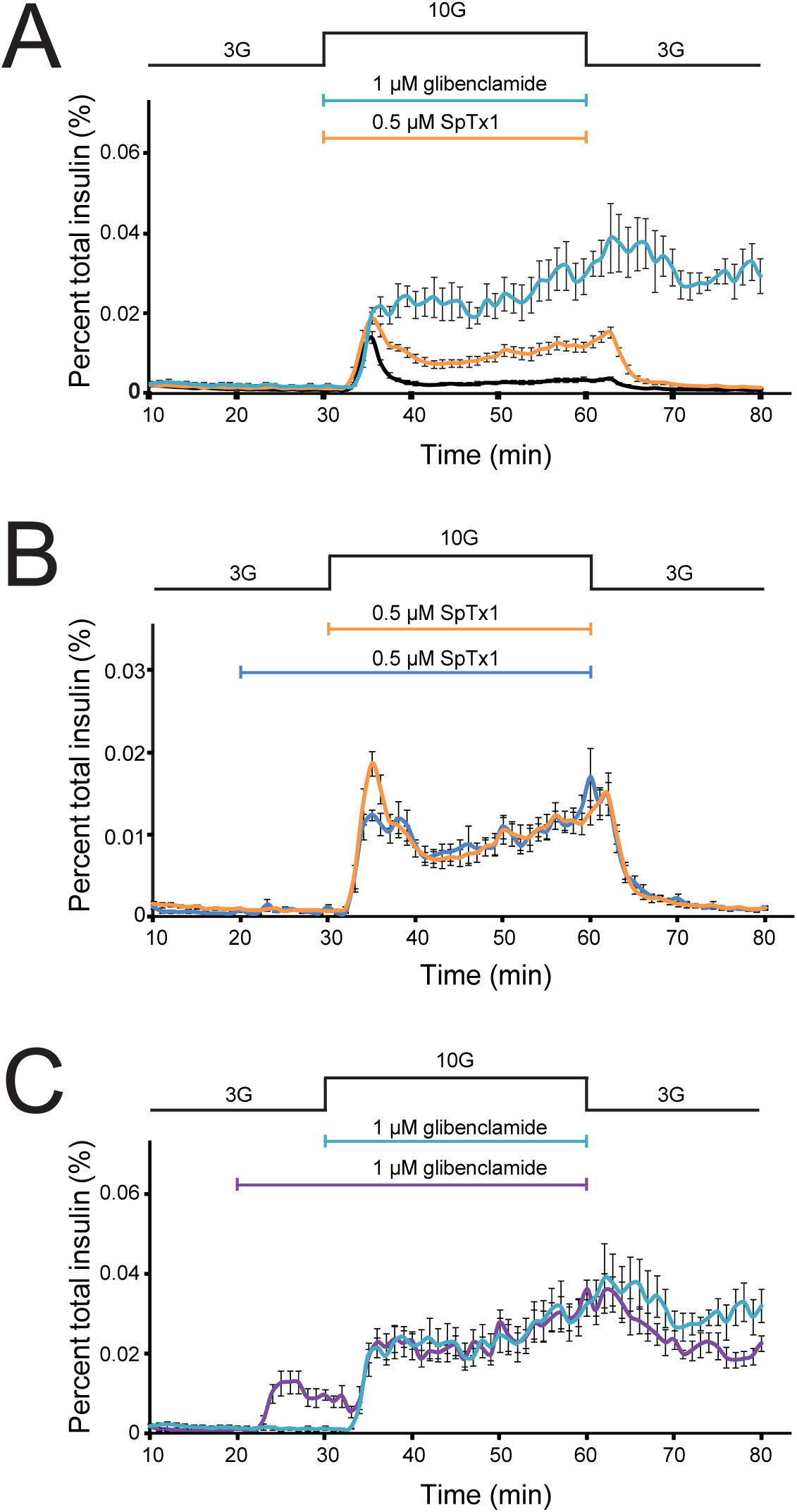
SpTx1 potentiates 10 mM glucose-stimulated insulin section from pancreatic islets of the ^Endo^mKir6.2^V108E^ mice. In all graphs, the amount of insulin per mL perifusion solution released from ∼180 islets from ^Endo^mKir6.2^V108E^ mice (8 - 12 weeks of age) as a percentage of the total insulin content is plotted against time. The rate of perifusion was 1 mL/min. All data points are plotted as mean ± SEM (*n* = 4 - 6), connected by lines color-coded for various conditions. (**A-C**) Insulin secretion from isolated islets of ^Endo^mKir6.2^V108E^ mice where the elevation of the glucose concentration in the perifusion solution from 3 mM to 10 mM and the return to 3 mM was as indicated by the black schematic outline at the top. In **A**, the cyan, orange, or black data plot represents the insulin secretion profile with the inclusion of 1 μM glibenclamide, 0.5 μM SpTx1, or neither; in **B**, the blue or orange data plot represents the insulin secretion profile with the inclusion of 0.5 μM of SpTx1 either 10 min prior to or at the time of raising the glucose concentration; and in **C**, the purple or cyan data plot represents the insulin secretion profile with the inclusion of 1 μM of glibenclamide either 10 min prior to or at the time of raising the glucose concentration. For ease of comparison, the orange data curve in **B** and cyan data curve in **C** are replotted from **A**.

Next, we tested whether SpTx1 could act as an effective secondary secretagogue to potentiate 10 mM glucose-stimulated insulin secretion (orange trace, Figure 9A). Because 0.2 to 1 μM SpTx1 did not potentiate insulin secretion from isolated wild-type mouse islets in 7 or 15 mM glucose (Figure 2), we first tested the effect of a concentration within this range on GSIS from isolated ^Endo^mKir6.2^V108E^ mouse islets. Inclusion of 0.5 μM SpTx1 in the perifusion solution potentiated insulin secretion in 10 mM glucose to a level between that in 10 mM glucose alone and 10 mM glucose plus glibenclamide. Following removal of high glucose stimulation by subsequent perifusion with 3 mM glucose, which also washed out extracellularly bound SpTx1, insulin secretion returned to the baseline level. This potentiating effect of SpTx1 on GSIS from islets of ^Endo^mKir6.2^V108E^ mice, not from those of wild-type mice, strongly implies that SpTx1 generated this effect by blocking Kir6.2 channels harboring the introduced V108E mutation.

We wondered whether SpTx1 diffused slowly into islets to reach β cells, and a pre-incubation of islets with SpTx1 would thus produce a greater potentiation of GSIS. To examine this possibility, we added SpTx1 10 minutes before raising glucose concentration to allow more time for equilibration. We found that, whether added earlier or at the same time as raising the concentration of glucose, SpTx1 had comparable potentiating effects on 10 mM glucose-stimulated insulin secretion (blue or orange trace, Figure 9B). The potentiating effects of glibenclamide were also comparable regardless of whether it was applied concurrent to or before glucose elevation (cyan or purple trace, Figure 9C). Adding these two inhibitors 10 minutes before the high glucose challenge also provided us an opportunity to evaluate the compounds as primary secretagogues, triggering insulin secretion in the presence of a non-stimulating concentration of glucose. In the presence of non-stimulating 3 mM glucose, glibenclamide triggered sizable secretion of insulin (purple trace, Figure 9C) whereas SpTx1 caused no detectable secretion of insulin above the baseline level (blue trace, Figure 9B).

Furthermore, we also examined the effect of SpTx1 on insulin secretion in the presence of 16.7 mM glucose, which is an empirically determined optimal concentration commonly used to stimulate a very high level of insulin secretion from mouse islets (Alcazar and Buchwald, 2019; Cerasi, 1975; Malaisse et al., 1967). Either glibenclamide or SpTx1 was added into the perifusion solution containing 3 mM glucose, 10 minutes before raising glucose to 16.7 mM. In the presence of 3 mM glucose, glibenclamide again stimulated sizable secretion of insulin, which decreased somewhat with time (purple trace, Figure 10). Raising glucose to 16.7 mM in the presence of glibenclamide caused much greater secretion of insulin than it did in the absence of glibenclamide (black trace versus purple trace, Figure 10). The same procedure was repeated with SpTx1. In the presence of 3 mM glucose, 0.5 μM SpTx1 did not stimulate observable secretion of insulin but did markedly potentiate insulin secretion in the presence of 16.7 mM glucose (blue trace), albeit less than that potentiated by 1 μM glibenclamide (purple trace, Figure 10). As we increased the SpTx1 concentration to 2 μM (vermilion trace), SpTx1 potentiated insulin secretion in the presence of 16.7 mM glucose to a level comparable to that caused by 1 μM glibenclamide (Figure 10). These observations establish a pair of practically equivalent concentrations for the two inhibitors to act as secondary insulin secretagogues. [For reference, both 2 μM SpTx1 (K_d_ = 20 nM; Figure 7C) and 1 μM glibenclamide are expected to inhibit 99% of the K^+^ current carried by their target channels on the cell surface (Proks et al., 2013).] However, at these equivalent concentrations, glibenclamide and SpTx1 did not have comparable abilities as primary secretagogues; SpTx1 stimulated much less insulin secretion than glibenclamide did as evidenced by their differing effects in non-stimulating 3 mM glucose (vermilion trace versus purple trace, Figure 10). These comparable secondary-secretagogue effects but differential primary-secretagogue effects between glibenclamide and SpTx1 are more clearly shown by the ratio of the mean insulin secretion in the presence of 1 μM glibenclamide to that in the presence of 2 μM SpTx1 (orange curve in the lower panel, Figure 10). Such a difference in the ratio was even more pronounced when 1 μM glibenclamide and 0.5 μM SpTx1 were compared (cyan curve in the lower panel, Figure 10).

**Figure 10.**
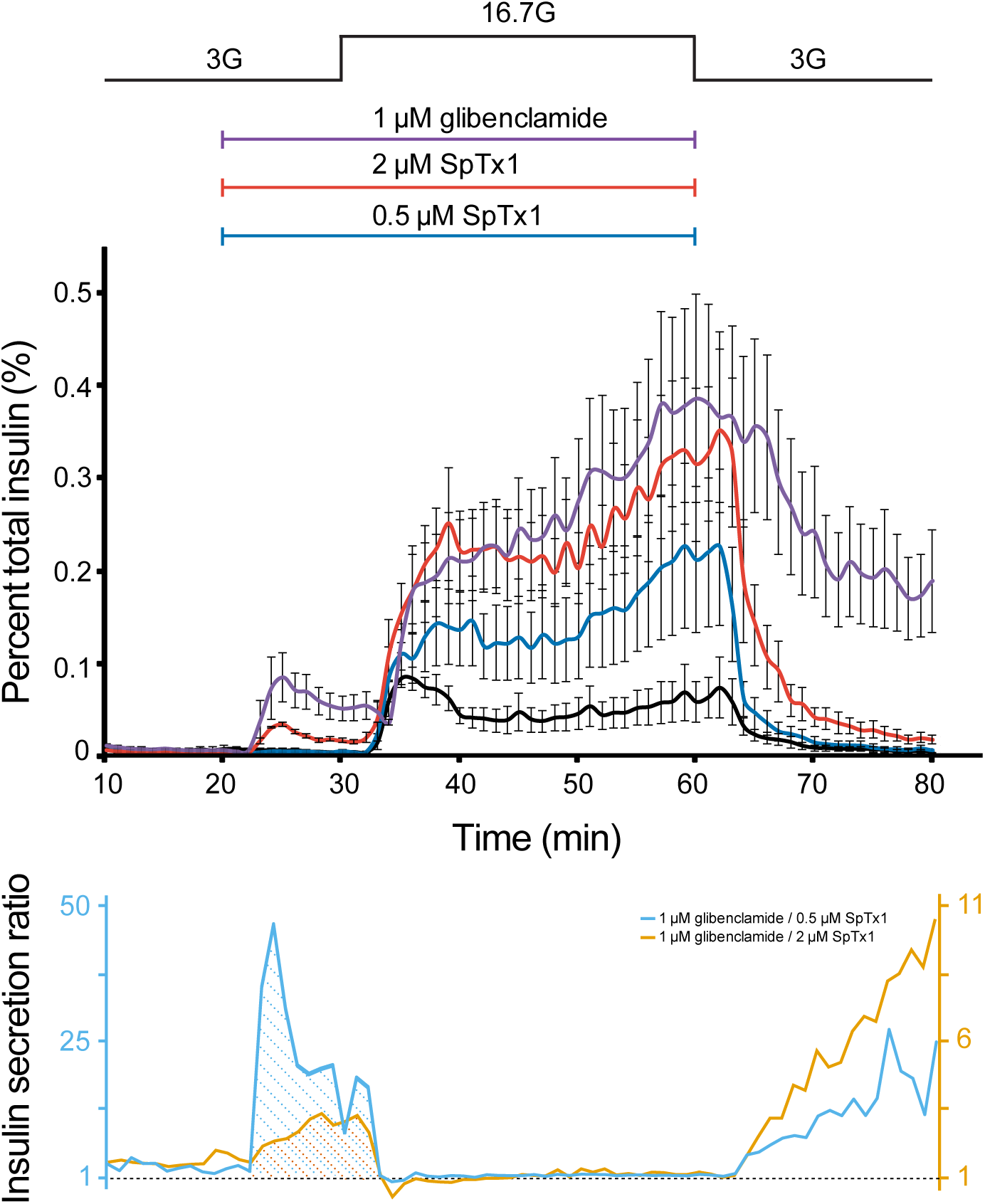
SpTx1 potentiates 16.7 mM glucose-stimulated insulin secretion from isolated pancreatic islets of ^Endo^mKir6.2^V108E^ mice. The amount of insulin per mL perifusion solution released from ∼180 islets from ^Endo^mKir6.2^V108E^ mice (8 - 12 weeks of age) as a percentage of their total insulin content is plotted against time. The rate of perifusion was 1 mL/min. All data points are plotted as mean ± SEM (*n* = 4), connected by lines color-coded for various conditions. The elevation of the glucose concentration in the perifusion solution from 3 mM to 16.7 mM and the return to 3 mM was as indicated by the black schematic outline at the top. The purple, vermilion, blue, and black data plots represent the insulin secretion profile with the inclusion of 1 μM glibenclamide, 2 μM SpTx1, 0.5 μM SpTx1, or neither inhibitor. The orange and cyan curves shown below the insulin secretion profiles represent the ratio between the amount of insulin secreted in the presence of 1 μM glibenclamide and that in the presence of 0.5 μM SpTx1 (cyan) or 2 μM SpTx1 (orange).

### SpTx1 counteracts the antagonistic effect of diazoxide on GSIS from pancreatic islets of the V108E-mutant mice

If SpTx1 acts as an effective secondary secretagogue by blocking the V108E-mutant mKir6.2, then SpTx1 is expected to counteract the effect of diazoxide, a so-called opener of K_ATP_ channels. In a static assay, 16.7 mM glucose stimulated insulin secretion from isolated ^Endo^mKir6.2^V108E^ mouse islets to a level above the baseline level in 3 mM glucose (Figure 11). Furthermore, 100 µM diazoxide lowered the detected signal in 16.7 mM glucose to the baseline level. SpTx1 at 1 µM, but not 0.02 µM, could effectively boost GSIS in the presence of diazoxide. In contrast, in the presence of non-stimulating 3 mM glucose, the levels of insulin secretion were comparably low under all four compared conditions.

**Figure 11.**
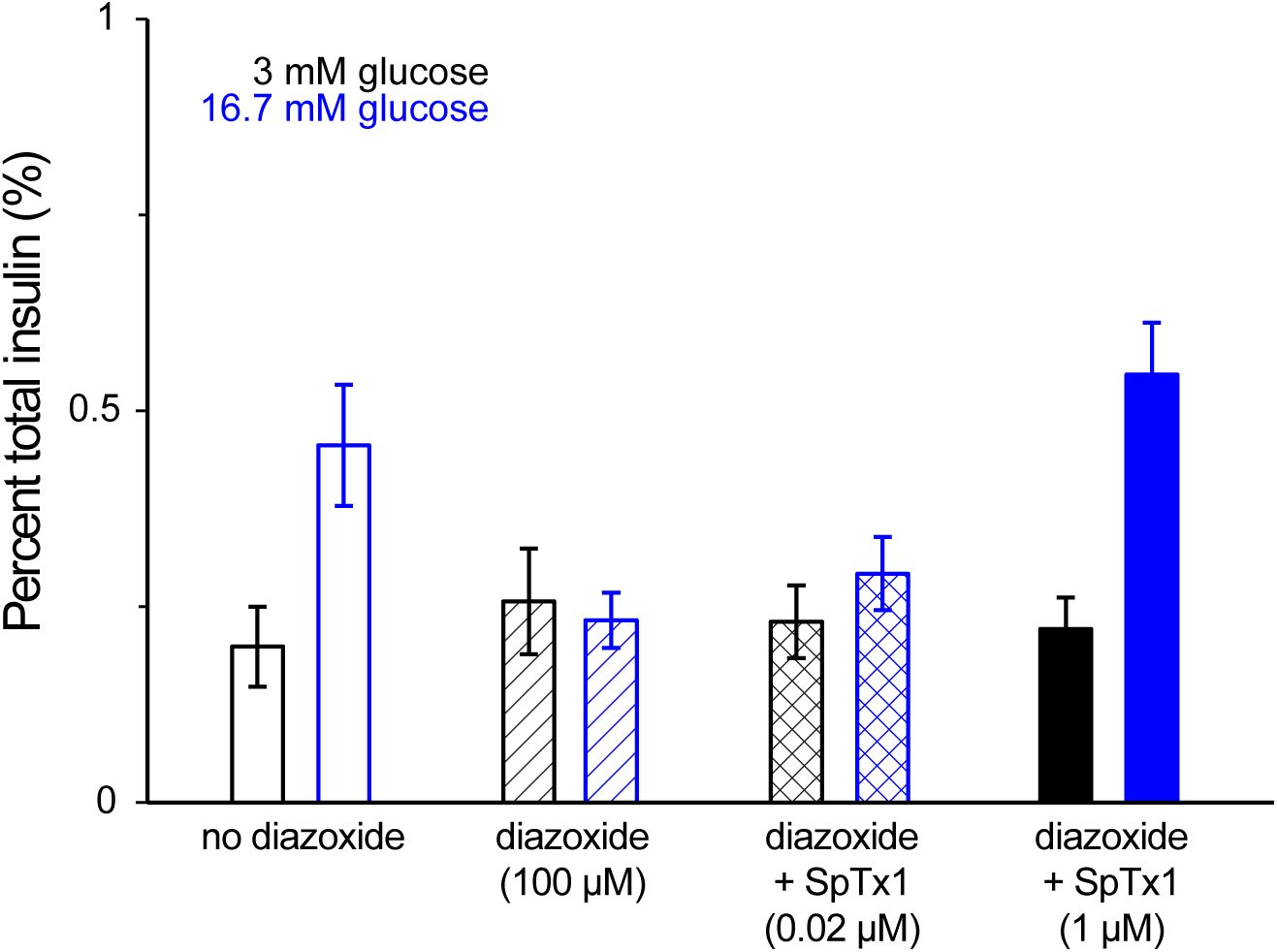
SpTx1 counteracts the antagonistic effect of diazoxide effect on GSIS from isolated pancreatic islets of ^Endo^mKir6.2^V108E^ mice. For each of the six independent experiments for a given condition as indicated, individual groups of 5 or 10 islets from ^Endo^mKir6.2^V108E^ mice (8 - 12 weeks of age) were incubated in the wells of a microwell plate. The secreted insulin as a percentage of the total insulin content of the islets (mean ± SEM; *n* = 6 for each case) is presented as histograms where black or blue colors code for the presence of 3 mM or 16.7 mM glucose in the bathing medium. The fill pattern inside each pair of rectangles represents a tested condition: glucose only (open), glucose + 100 μM diazoxide (diagonal lines), glucose + 100 μM diazoxide + 0.02 μM SpTx1 (crossed lines), and glucose + 100 μM diazoxide + 1 μM SpTx1 (filled). For the two key group comparisons, the P value of two-tailed student’s *t*-test is 0.028 between the glucose only group and the glucose + 100 μM diazoxide group and is 0.006 between the glucose + 100 μM diazoxide group and the glucose + 100 μM diazoxide + 1 μM SpTx1 group, all in the presence of 16.7 mM glucose.

### SpTx1 increases the plasma insulin level and lowers the blood glucose concentration in diabetic mice

Our ability to confer high SpTx1 sensitivity on mKir6.2 in mice enabled us to examine whether SpTx1 can actually trigger impactful insulin secretion and thus lower the elevated blood glucose level in diabetic mice that overexpress mKir6.2 activity (Figure 12). We started with an NDM-model diabetic-mouse line previously constructed by the Nichols’ group, which contained an inducible mutant mKir6.2 transgene in the *Rosa26* locus, hereafter dubbed ^Rosa26^mKir6.2^NDM^ (Remedi et al., 2009). After a 9-day induction to overexpress the constitutively active mutant mKir6.2 in the pancreas of ^Rosa26^mKir6.2^NDM^ mice, the overnight-fasted blood glucose level of the mice remained highly elevated during the 2-hour observation period (Figure. 12A). We found that an IV injection of SpTx1 (1 mg/kg) or its vehicle saline had no markedly meaningful effect on the elevated blood glucose levels of diabetic ^Rosa26^mKir6.2^NDM^ mice and did not trigger a rise of insulin in the plasma of their circulating blood collected during the observation period (filled or open triangles, Figure 12A and 12C). These findings are expected because mKir6.2 exhibits very low affinity for SpTx1.

**Figure 12.**
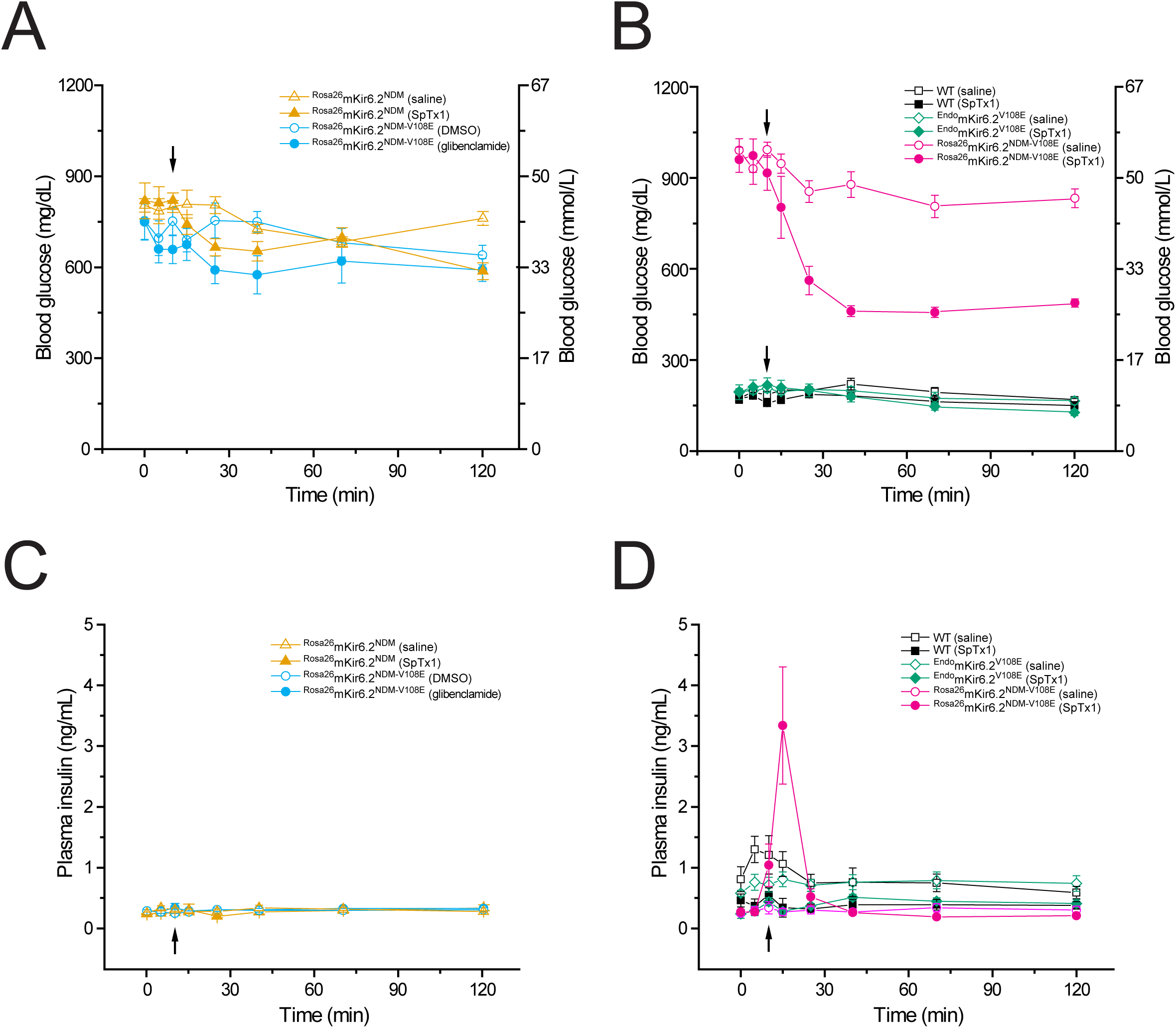
SpTx1 triggers a transient rise of plasma insulin and lowers the elevated blood glucose level and in diabetic ^Rosa26^mKir6.2^NDM-V108E^ mice. (**A-D**) Blood glucose (**A** and **B**) and corresponding plasma insulin (**C** and **D**) levels (mean ± SEM) of overnight fasted mice (8 - 12 weeks of age) at indicated time points during a 2-hour observation period. (**A** and **C**) SpTx1 (1 mg/kg, filled orange triangles) or its vehicle saline (open orange triangles) was intravenously administered in ^Rosa26^mKir6.2^NDM^ mice (*n* = 5 for each case) and glibenclamide (40 mg/kg, filled cyan circles) or its vehicle DMSO (open cyan circles) was intraperitoneally administered in ^Rosa26^mKir6.2^NDM-V108E^ mice (*n* = 5 each) as indicated by the arrow. (**B** and **D**) SpTx1 (1 mg/kg, filled symbols) or its vehicle saline (open symbols) was intravenously administered in wild-type (black squares, *n* = 5 each), ^Endo^mKir6.2^V108E^ (green diamonds, *n* = 5 each), and ^Rosa26^mKir6.2^NDM-V108E^ (magenta circles, *n* = 10 each) mice as indicated by the arrow. For the comparison of the plasma insulin levels of ^Rosa26^mKir6.2^NDM-V108E^ mice at 5 minutes post administration, the P value of two-tailed student’s *t*-test is 0.005 between the vehicle group and the SpTx1 group.

To increase the affinity of mKir6.2 in ^Rosa26^mKir6.2^NDM^ mice for SpTx1, we then introduced the V108E mutation into all endogenous and transgenic copies of mKir6.2 within their genome (Methods). The resulting mice are denoted as ^Rosa26^mKir6.2^NDM-V108E^. Like the original ^Rosa26^mKir6.2^NDM^ mice, after a 9-day induction to overexpress the mutant mKir6.2 in the pancreas of ^Rosa26^mKir6.2^NDM-V108E^ mice, their overnight-fasted blood glucose level also became highly elevated in a sustained manner. A single bolus application of glibenclamide (40 mg/kg) or its vehicle DMSO neither markedly lowered the elevated blood glucose level (filled or open circles, Figure 12A) nor caused a rise of insulin in the plasma (circles, Figure 12C) of diabetic ^Rosa26^mKir6.2^NDM-V108E^ mice during the observation period. This observation is consistent with the ineffectiveness of acute glibenclamide treatments reported in the original study of ^Rosa26^mKir6.2^NDM^ mice (Remedi et al., 2009). Thus, in the present context, ^Rosa26^mKir6.2^NDM-V108E^ mice retained the basic characteristics of inducible diabetic ^Rosa26^mKir6.2^NDM^ mice.

Next, we tested whether SpTx1 could influence the blood glucose level of ^Rosa26^mKir6.2^NDM-V108E^ mice. Following an IV injection of SpTx1 (1 mg/kg) but not saline, the elevated blood glucose level of diabetic ^Rosa26^mKir6.2^NDM-V108E^ mice dropped markedly over 30 minutes and this lower level of blood glucose persisted over the remaining observation period (filled or open magenta circles, Figure 12B). In contrast, either SpTx1 or saline (filled or open symbols) had little effects on the blood glucose levels of wild-type and non-diabetic ^Endo^mKir6.2^V108E^ mice (squares and diamonds) for the observed period (Figure 12B). Moreover, the IV injection of SpTx1, but not of saline, triggered a transient but sizable rise of insulin in the plasma of diabetic ^Rosa26^mKir6.2^NDM-V108E^ mice, but not in that of wild-type or non-diabetic ^Endo^mKir6.2^V108E^ mice (Figure 12D). These results show that SpTx1 promotes transient insulin secretion *in vivo* and markedly lowers the highly elevated blood glucose level of diabetic ^Rosa26^mKir6.2^NDM-V108E^ mice.

## Discussion

We originally discovered SpTx1 on the basis of its potent inhibition of hKi6.2 with a K_d_ of 15 nM. In the presence study, we found that at 0.2 µM concentration, SpTx1 failed to depolarize β cells, and at a concentration as high as 1 μM, SpTx1 had no observable effects on GSIS from β cells in isolated pancreatic islets of wild-type mice (Figures 2 and 3). This ineffectiveness led us to determine that SpTx1 blocks mKir6.2 with much lower potency than it blocks hKir6.2 (Figure 4). Through a series of mutagenesis studies, we found that this low-affinity interaction between mKir6.2 and SpTx1 stems from the presence of a valine residue at position 108 in mKir6.2 whereas it is a glutamate residue in hKir6.2 (Figure 5). Replacing the valine residue in mKir6.2 by a glutamate residue confers the high affinity of hKir6.2 for SpTx1 upon mKir6.2 (Figures 6 and 7). This mechanistic information led us to create the SpTx1-sensitive ^Endo^mKir6.2^V108E^ transgenic mouse line. Using isolated islets of these transgenic mice, we found that contrary to the expectation of Kir6.2 inhibition causing sufficient V_m_ depolarization and robust insulin secretion, SpTx1 does not impactfully depolarize β cells in glucose concentrations less than 5 mM nor act as an effective primary insulin secretagogue in 3 mM glucose. Nonetheless, SpTx1 depolarizes β cells and elicits action potentials in higher glucose and acts as an effective secondary secretagogue, strongly potentiating GSIS from these islets (Figures 8 and 9).

SpTx1 inhibits K_ATP_ channels by blocking their ion-conduction pore whereas glibenclamide inhibits the channels by disrupting their nucleotide-dependent gating. Despite this difference, if both SpTx1 and glibenclamide promote insulin secretion solely by inhibiting K^+^ currents through mKir6.2, increasing the concentration of SpTx1 to a sufficiently high level should produce effects that would match those produced by glibenclamide, in the presence of whether a non-stimulating or an optimal stimulating concentration of glucose. We observed that glibenclamide triggered sizable secretion of insulin in 3 mM glucose and potentiated insulin secretion in 16.7 mM glucose to a much higher level than glucose alone (Figure 10). In contrast, 0.5 μM SpTx1 did not trigger observable insulin secretion in 3 mM glucose nor potentiate insulin secretion in 16.7 mM glucose as strongly as glibenclamide. However, at 2 μM, SpTx1 did produce such a strong secondary secretagogue effect in 16.7 mM glucose that it potentiated GSIS to a level comparable to what was caused by glibenclamide, but it failed to act as a strong primary secretagogue to simulate as much insulin secretion in 3 mM glucose as glibenclamide did. Thus, at a sufficiently high concentration, SpTx1 can act as an effective secondary insulin secretagogue like glibenclamide but it does not appear to act as an effective primary insulin secretagogue.

One possible reason for these differing effects between SpTx1 and glibenclamide in a non-stimulating glucose concentration versus an optimal stimulating glucose concentration is that SpTx1 potentiated GSIS by acting on an off-target. However, we did not observe any meaningful potentiating effects of SpTx1 on GSIS from isolated islets of wild-type mice (Figure 2; see also Figures1B and 3) before we conferred a high SpTx1 affinity on their endogenous mKir6.2 (Figures 9 and 10; see also Figure 8). Furthermore, SpTx1 can counteract the inhibitory effect of the K_ATP_ opener diazoxide on GSIS (Figure 11). Thus, a potential off-target mechanism is unlikely.

An alternative possibility is that SpTx1 does not have all of the pharmacological actions of glibenclamide because glibenclamide may stimulate insulin secretion not solely by inhibiting Kir6.2 channels in the plasma membrane of β cells. Glibenclamide has been shown to cause an additional amount of insulin secretion, even after the membrane potential was clamped to a depolarized level by raising the extracellular K^+^ to a constant 30 mM such that the impact of any alteration of the Kir6.2 activity on the membrane potential of the β cells and consequently the Ca_v_ activity would become negligible (Geng et al., 2007). Thus, glibenclamide may have an additional effect downstream of the cell surface K_ATP_ channels, and consequently the extracellular Kir6.2-blocker SpTx1 cannot effectively mimic every action of glibenclamide. Many studies have shown that glibenclamide interacts with other proteins including syntaxin-1A and Epac2 (Eliasson et al., 1996; Hinke, 2009; Kang et al., 2011; Lehtihet et al., 2003; Renstrom et al., 2002; Shibasaki et al., 2014; Tian et al., 1998; Zhang et al., 2009). In principle, glibenclamide may also act on some intracellular K_ATP_ channels that SpTx1 cannot access (Geng et al., 2003).

Regarding the application of SpTx1 in *in vivo* studies, on one hand, SpTx1’s characteristics, which allow the toxin to function only as an effective secondary but not as an effective primary secretagogue, make it unsuitable when a strong primary secretagogue effect is needed. On the other hand, being an effective secondary but not an effective primary secretagogue, SpTx1 will only strongly promote insulin secretion when blood glucose level becomes highly elevated but it will not meaningfully stimulate insulin secretion at a resting level of blood glucose – a desirable outcome unlikely to cause unwanted hyperinsulinemia and hypoglycemia (Asplund et al., 1983). Consistent with these predictions, we observed that the administration of SpTx1 did not trigger insulin secretion nor lower the blood glucose levels in non-diabetic ^Endo^mKir6.2^V108E^ mice, but it caused transient insulin secretion and markedly lowered the highly elevated glucose level in diabetic ^Rosa26^mKir6.2^NDM-V108E^ mice (Figure 12). In the future, it will be interesting to learn in animal models whether the transient characteristic of insulin secretion caused by SpTx1 helps to mitigate problems associated with over-stimulation of insulin secretion and “exhaustion” of β cells, which can occur with the use of sulfonylureas (Maedler et al., 2005; Matthews et al., 1998).

As experimental tools, sulfonylureas and SpTx1, while both inhibiting K_ATP_ channels, have important and different characteristics. The pancreatic K_ATP_ channels made of Kir6.2 and SUR1 are inhibited by ATP and activated by MgADP (Gribble and Reimann, 2003; Proks et al., 2013). The stimulatory effect of MgADP is antagonized by binding of sulfonylureas to SUR1. This apparent inhibitory effect, together with the ATP-mediated inhibition, decreases the channel open probability (P_o_) to near zero. However, without ATP, sulfonylureas alone can only inhibit about 2/3 of the channel current. Thus, the inhibitory effect of sulfonylureas depends on the cellular metabolic state. In contrast, the inhibitory effect of SpTx1 on the Kir6.2 pore in the K_ATP_ channel complex should not depend on the cellular metabolic state. This expectation stems from the observation that SpTx1 blocks currents through a constitutively active Kir6.2 mutant and those through K_ATP_ channels activated with azide that lowers the concentration of intracellular ATP (Figures 4-7) comparably well. However, SpTx1 is of no use when the Kir6.2 pore in the K_ATP_ protein complex is inherently insensitive or is experimentally rendered insensitive to SpTx1, such as that of the wild-type mice and those carrying natural or engineered SpTx1-resistant mutations. Sulfonylureas also have limitations in inhibition K_ATP_ channels. Mutations in SUR1 could affect K_ATP_ channels such that their current is not adequately inhibited by sulfonylureas (Proks et al., 2004). The K_ATP_ complexes containing SUR2 have apparently low sulfonylurea sensitivity, e.g., in cardiac and some neuronal K_ATP_ channels (Nichols, 2016; Seino and Miki, 2003). Another important difference between the two inhibitors is that SpTx1 is not membrane-permeable and acts from the extracellular side whereas sulfonylureas are membrane-permeable and lodge into the transmembrane segments of SUR proteins (Lee et al., 2017; Li et al., 2017; Martin et al., 2017; Schatz et al., 1977; Wu et al., 2018).

Because SpTx1 and sulfonylureas have different characteristics, these inhibitors may have different therapeutic benefits. For example, sulfonylureas have only limited effects on the neurological disorders of patients with DEND (Hattersley and Ashcroft, 2005; Pipatpolkai et al., 2020). To this end, it will be interesting to test whether SpTx1 can help to mitigate these neurological disorders in animal models by directly delivering it into the cerebrospinal fluid.

In summary, we have conferred high-affinity SpTx1 inhibition upon mKir6.2 using the point-mutation V108E to mimic the human ortholog in a heterologous expression system, and introduced this mutation into mKir6.2 in wild-type and NDM-model mice via genome editing. Studying isolated pancreatic islets of SpTx1-sensitive knock-in mutant ^Endo^mKir6.2^V108E^ mice (without NDM-causing mutations), we have found that SpTx1 depolarizes V_m_, triggering action potentials, and that it acts as a secondary secretagogue that potentiates insulin secretion in the presence of high glucose, as effective as the sulfonylurea glibenclamide. However, in a low glucose condition, SpTx1 does not trigger action potentials nor acts as a primary secretagogue as effective as glibenclamide. Furthermore, an application of SpTx1 in ^Endo^mKir6.2^V108E^ mice neither triggers meaningful insulin secretion nor lowers their blood glucose from the resting level. By contrast, in SpTx1-sensitive and diabetic ^Rosa26^mKir6.2^NDM-V108E^ mice, an application of SpTx1 causes a transient rise of plasma insulin and markedly lowers their highly elevated blood glucose level. These features of the present experimental tool SpTx1 point to the potential therapeutic benefit of a Kir6.2 inhibitor with required pharmaceutical characteristics.

## Acknowledgements

We thank J. Bryan (Baylor College of Medicine) and L. Jan (University of California, San Francisco, CA) for sharing Kir6.2 and SUR1 cDNAs; C. Nichols (Washington University) for sharing the diabetic model mice; and the University of Pennsylvania Transgenic and Chimeric Mouse Facility for the transgenic mouse service and the Diabetes Research Center Core for performing the perifusion and RIA assays of mouse pancreatic islet β cells (P30-DK19525). The present study in the early phase was supported by the National Institute of Diabetes and Digestive and Kidney Diseases (DK109979) to Z.L., and in the later phase that did not directly involve mice by the National Institute of General Medical Sciences (GM55560) to Z.L. T.H. was in part supported by DK098517.

## Author contributions

Z.L. conceived the study; Y. Ramu, J. Yamakaze, Y. Zhou, T. Hoshi and Z. Lu designed the study; Y. Ramu, J. Yamakaze, Y. Zhou and T. Hoshi performed the experiments; Y. Ramu, J. Yamakaze, Y. Zhou, T. Hoshi and Z. Lu analyzed and interpreted the data and wrote the manuscript.

## Data availability

The datasets used in the present study are included in the Supplementary Materials.

## Methods

### Ethics and approval for animal experiments

The Institutional Animal Care and Use Committee at the University of Pennsylvania reviewed and approved this study (Protocol Numbers: 804489 and 804811). All animal welfare considerations were taken and the study was performed in strict accordance with the recommendations in the Guide for the Care and Use of Laboratory Animals of the National Institutes of Health. All animal care was provided by experienced staff and researchers, and surgical procedures were performed after euthanasia by researchers who were trained at the University Laboratory Animal Resources of the University of Pennsylvania. Every effort was made to minimize suffering and distressed of mice. The studies were performed in compliance with the ARRIVE guidelines (https://arriveguidelines.org/).

### Mutagenesis and electrophysiological recordings

All mutant Kir6.2 cDNAs were produced through PCR-based mutagenesis and confirmed by DNA sequencing. The Kir6.2 and SUR1 cRNAs were synthesized with T7 polymerase using the corresponding linearized cDNAs as templates. A two-electrode voltage-clamp amplifier (Oocyte Clamp OC-725C, Warner Instruments Corp.) was used to record channel currents from *Xenopus* oocytes injected with cRNA encoding specific wild-type or mutant Kir6.2 channels, with or without co-injection with cRNA encoding SUR1. All recordings were performed at room temperature. To activate K_ATP_ channel currents, 3 mM azide was added to the bath solution. The resistance of electrodes filled with 3 M KCl was 0.2 - 0.4 MΩ. To elicit currents through the channels, the membrane potential of oocytes was stepped from the holding potential of 0 mV to - 80 mV then to +80 mV before returning back to 0 mV. The bath solution contained (in mM): 100 KCl, 0.3 CaCl_2_, 1.0 MgCl_2_, and 10 HEPES, and BSA (50 μg/mL), where pH was titrated to 7.6 with KOH. Recombinant SpTx1, produced as previously described (Ramu et al., 2018), was added to the bath solution in concentrations as specified in relevant figures. The range for the number of measurements was determined on basis of previous studies (Ramu et al., 2018; Ramu and Lu, 2019). The variations in data values reflected both biological variability and technical errors. No experiments were excluded.

Electrophysiological membrane potential (V_m_) measurements from individual cells in isolated but intact islets cultured on glass coverslips were performed using the perforated whole-cell patch-clamp method as described (Yang et al., 2021). Wax-coated electrodes (G85150T, Warner) were back-filled with the intracellular solution containing (in mM): 76 K_2_SO4, 10 KCl, 10 NaCl, 6 MgCl_2_, 30 mannitol, 30 HEPES, where pH was titrated to 7.2 at 35 °C with *N*-methyl-*D*-glucamine (NMG). The tip of the pipette was filled with this solution and the remainder was back filled with the same solution plus β-escin (6 or 8 µM). The free Mg^2+^ concentration of this solution, taking the divalent cation-chelating action of SO_4_^2–^ into consideration, is estimated to be about 2 mM. The extracellular solution contained (in mM): 135 NaCl, 4 KCl, 2 CaCl_2_, 2 MgCl_2_, 10 HEPES, where pH was titrated to 7.4 at 35 °C with NMG. Glucose was added to this solution as required and the osmolarity was adjusted to ∼300 mOsm with mannitol. All salts were from Millipore-Sigma. Using the solutions described above, the initial input resistance of the electrode was typically 4 to 8 MΩ. The electrophysiological recordings were performed under a continuous perfusion condition, 0.3 to 0.4 mL/min (Instech), at ∼35 °C (TC-124A and TC-344B Warner). Islets were equilibrated in the recording chamber for 10 to 15 min before measurements were made. Adequate whole-cell access was achieved typically within 5 to 10 min. SpTx1 was applied with 0.3% w/v BSA.

The output of the patch-clamp amplifier (Axopatch 200B, Molecular Devices) was digitized at 2 kHz and stored for later analysis using Igor Pro (v8 or v9, Wavemetrics). The liquid junction potential has been subtracted from the results. To estimate the average V_m_ values under different conditions, the V_m_ traces were digitally Gaussian filtered at 0.1 Hz and the filtered data points in the last 50 s of a segment of interest were averaged. The filtered traces are superimposed on the illustrative original traces in the figures where appropriate. The variations in data values reflected both biological variability and technical errors (Figures 3 and 8, right panels). No data, which were successfully recorded over sufficient durations that allowed for adequate examination the membrane potential under the various specified conditions, were excluded.

### Generation of ^Endo^mKir6.2^V108E^ and ^Rosa26^mKir6.2^NDM-V108E^ transgenic mice

To confer the Kir6.2-like high SpTx1-sensitivity on endogenous mKir6.2 channels in wild-type mice, we employed the CRISPR-Cas9 genome editing technique (Ran et al., 2013). The sequences of *in vitro* transcription template of a guide RNA (gRNA) and the single-stranded oligonucleotide (ssODN) donor templates are provided below.

Synthesized DNA templates (IDT Technologies) for gRNA targeting the Kir6.2 DNA sequence (Genbank, Accession Number NC_000073.6) also contained a T7 promoter sequence. The sequence of the DNA template for the gRNA from the 5’ end to the 3’ was: <u>GAAATTAATACGACTCACTATAGGGAGA</u>**GCTTGTGACGCAGGGCACAT***GTTTTAGA GCTAGAAATAGCAAGTTAAAATAAGGCTAGTCCGTTATCAACTTGAAAAAGTGGCACCGAGTCGGTGCTTTTTT* where the T7 promoter sequences are underlined, target DNA sequences bolded, and tracrRNA italicized; whereas that for the ssODN donor template was: GGACCTCGATGGAGAAAAGGAAGGCAGATGAAAAGGAGTGGATGCTTGTGACGCA <u>GGGC**T**C</u>*G*TT*A*GTGCCCTCTCCGGGGGCCAGGTCACCGTGGGCGAAGGCGATGAGCC ACCAGACCATGGCAAAGA where the sequence of the restriction enzyme (BanII) cleavage site is underlined, the intended non-synonymous substitution is bolded and synonymous ones italicized (Figure S1C).

*In vitro* transcription from the gRNA template or a Cas9 plasmid (T7-Cas9-HA-2NLS) was performed to produce gRNA or Cas9 mRNA using mMESSAGE mMACHINE T7 ULTRA kit (Invitrogen, AM1344). The ssODN stock solution was prepared according to manufacturer’s recommendation (IDT Technologies). The freshly prepared solution, which contained 100 ng/µL Cas9 mRNA, 100 ng/µL gRNA, 200 ng/µL ssODN, 0.1 mM EDTA, and 10 mM Tris titrated to pH 7.5, was injected into cytoplasm of FVB mouse embryos at the Transgenic and Chimeric Mouse Facility (TCMF) of the University of Pennsylvania Perelman School of Medicine (UPENN-PSOM). The injected embryos were implanted into the uterus of pre-prepared surrogate female mice for obtaining germline-transmitting founder ^Endo^mKir6.2^V108E^ mutant mice. The intended T323A mutation we designed and the 5 adjacent nucleotides in the DNA created a cleave site for the restriction enzyme BanII (underline, Figure S1A-S1C), which was used to assist the identification of genotypes.

Double transgenic ^Rosa26^mKir6.2^NDM^ mice on the C57BL/6J (B6J) background were obtained from the laboratory of Dr. Colin Nichols (Washington University, St. Louis, MO)(Remedi et al., 2009). This mutant mouse line contained two unlinked transgenes. One encodes for a tamoxifen-dependent, pancreas-specific Cre recombinase (Pdx-*Cre*^ERT2^) and the other for a Cre-inducible copy of mutant Kir6.2 gene at the *Rosa26* locus, in addition to the wild-type Kir6.2 gene (*Kcnj11*) at the endogenous locus. The translation of mutant Kir6.2 transgene would harbor the K185Q mutation and lack N-terminal 30 residues (ΔN30). By crossing their ^Rosa26^mKir6.2^NDM^ mice with our mutant ^Endo^mKir6.2^V108E^ mice, we created a hybrid mutant mouse line on the B6.FVB background whose genome contained the two unlinked transgenes and the mutated endogenous Kir6.2 gene with homozygous V108E mutation. Furthermore, using the hybrid mutant line and the CRISPR-Cas9 targeting strategy described above, we introduced the V108E mutation into the mutant Kir6.2 transgene at the *Rosa26* locus, and the resulting germline-transmitting mutant mice are dubbed as ^Rosa26^mKir6.2^NDM-V108E^, in which only the mutant Kir6.2 transgene contain K185Q and ΔN30 mutations but both transgenic and endogenous Kir6.2 genes contain V108E mutation. All mice used in the study were subsequently generated in-house and maintained in a UPENN-PSOM animal facility.

### Mice maintenance and their humane endpoints

All mice were housed in ventilated, sterilized polysulfone mouse cages (AN75, Ancare Corp., Bellmore, New York, USA) containing Bed-o’Cobs absorbent bedding (Andersons Lab Bedding, Maumee, Ohio, USA). Irradiated LabDiet 5053 feed (PicoLab Rodent Diet 20, Fort Worth, Texas, USA) and autoclaved acidified-water (pH 2.5 - 2.8) were provided *ad libitum*. Shepherd’s shack and nesting material were also provided in the cages to minimize distress of mice. Cages were changed weekly in a Class A2 biological safety cabinet (Baker SG403A-ats, Sanford, Maine, USA). The air in the mouse-holding room was exchanged with filtered air at a rate of 15 times of room volume per hour. Room temperature was maintained between 20 and 25°C, and relative humidity between 30 and 70%. Lighting of the room was on and off alternately every 12 hours where the light phase was from 7:00AM to 7:00PM. A vivarium-wide rodent health monitoring program showed the absence of the following pathogens: MHV, Sendai virus, MVM, MPV1/2, TMEV, PVM, Reo-3, EDIM, Ectromelia virus, LCMV, MAdV, K-Virus, Polyomavirus, MCMV, Mouse Thymic Virus, Hantavirus, *Mycoplasma pulmonis, Citrobacter rodentium, Clostridium piliforme, Corynebacterioum kutscheri*, CAR *bacillus.*, *Salmonella* sp.*, Klebsiella pneumoniae, Streptococcus pneumoniae, Streptobacillus moniliformis, Encephalitozoon cuniculi, Myobia musculi, Myocoptes musculinus, Radfordia affinis, Aspiculuris tetraptera, Syphacia obvelata, Giardia muris*.

The animal cohort in this study included wild-type and mutant mice at 8-12 weeks of age. The designed end point of the study was defined as individual mice reaching that age range and euthanized for various types of experiments. We monitored the general health condition and behavior of each mouse at least 3 times per week during maintenance and continuously during the 2-hour observation experiment. If a mouse was suspected to have clinical signs of pain and distress, this mouse was then monitored and checked one or several times daily. We used the following criteria to determine the humane endpoints at which mice of any age were euthanized within one hour upon our determination: clinical signs of pain and distress, such as hunched posture, inactivity, dehydration, abdominal enlargement caused by intestine obstruction, increased respiratory effort manifested as increased intercostal or subdiaphragmatic retraction and gasping or breathing with an open mouth, raffled fur coat, emaciated body condition (e.g., weight loss of >20%), severe lethargy manifested as unwillingness to ambulate more than a few steps when gently stimulated with a gloved finger, or cold to the touch.

### Mouse genomic DNA isolation and genotyping

For genomic DNA isolation, a tail sample (∼2 mm) or a clipped toe from each mouse was digested at 55 ℃ overnight with gentle mixing in 0.5 mL solution containing 50 mM Tris titrated to pH 8.0, 100 mM EDTA, 0.5 % SDS, and 0.5 mg/mL Proteinase K. The high molecular weight genomic DNA in the digestion sample was extracted with a mixture of phenol and chloroform and precipitated using cold ethanol with 0.3 M sodium acetate titrated to pH 6.0. The precipitated genomic DNA was resuspended in 0.1 mL of a solution containing 1 mM EDTA and 10 mM Tris titrated to pH 8.0; 1 μL of which was used as the DNA template and added into a 25 μL PCR mixture containing 50 mM KCl, 1.5 mM MgCl_2_, 20 mM Tris triturated to pH 8.4, 0.2 mM dNTPs, 0.2 μM of each forward and reverse oligonucleotide primers described below, and 1 U of Platinum Taq DNA polymerase (Invitrogen, 10966034). PCR reactions took place in a thermal cycler (Mastercycler 5333; Eppendorf) with an initial denaturation step at 95 ℃ for 2 min and 30 amplification cycles (denaturation at 95 ℃ for 30 s, annealing at 60 ℃ for 30 s, and extension at 72 ℃ for 45 s for <0.7 kb or 150 s for ∼2 kb), and followed by a final 5 min extension step at 72 ℃. 5 μL of individual PCR products was subjected to electrophoresis on 1 % agarose gel, stained with 0.5 μg/mL ethidium bromide, and evaluated against a DNA ladder (Thermo Scientific, SM1331). The initial screen was performed on all resulting PCR products by the expected size (∼2 kb) and by their susceptibility to BanII restriction digestion.

By subsequent DNA sequencing of the whole ∼2 kb PCR product, we verified that the BanII-cut products contained the mutated codon for a glutamate residue at position 108 (Figure S1A and S1B), whereas the uncut products contained the wild-type codon for valine. Moreover, sequencing of the entire coding region of the Kir6.2 gene in the ^Endo^mKir6.2^V108E^ mice confirmed the two intended synonymous substitutions in the DNA triplets that encode amino acids threonine 106 and asparagine 107 without any other nucleotide changes. These intended substitutions were designed in the ssODN donor template to prevent re-editing by Cas9 while preserving their original amino acid identities. However, sequencing of the entire coding region of the mutant Kir6.2 transgene in the ^Rosa26^mKir6.2^NDM-V108E^ mice revealed a single unintended, non-synonymous substitution in the DNA codon that would lead to an additional N107Y mutation in the Kir6.2 protein (Figure S1C and S1D). Fortunately, this N107Y mutation had little effect on the affinity of channels for SpTx1 (Figure S2).

Specific primer pairs were designed using published genomic sequences for the mouse *Kcnj11* gene (Genbank accession: NC_000073.6) or the *Rosa26*locus (Genbank accession: NC_000072.6). Using Primer-BLAST (NIH) on default settings, each candidate pair was checked against the Refseq representative genomes of *Mus musculus*(taxid: 10090) to ensure a single expected PCR product. The oligonucleotide sequences of the synthesized primers (Sigma-Genosys) and the predicted sizes of the PCR products are given below:

1. For amplifying the entire coding region of the Kir6.2 gene at the endogenous locus, a 1,954 bp PCR product was expected with primers 5’-GGTAGACTTATCCCGCCGTG-3’ and 5’-TGGGGGCTCAGTAAGCAATG-3’
2. For amplifying the entire coding region of the Kir6.2 transgene at the *Rosa26* locus, a 1,981 bp PCR product was expected with primers 5’-GAGGCTACTGCTGACTCTCAA-3’ and 5’-GCTCGTCAAGAAGACAGGGC-3’
3. For testing the BanII-susceptibility, a 760 bp product was expected using PCR product from 1 or 2 as the template with primers 5’-CGCCCACAAGAACATTCGAG-3’ and 5’-GGTGATGCCCGTGGTTTCTA-3’. Upon exposure to BanII restriction enzyme, the 760 bp PCR product containing the designed V108E mutation will be cut into 565 bp and 195 bp fragments whereas the wild-type PCR product will remain uncut.
4. For verifying the presence of the Cre gene, a 429 bp PCR product was expected with primers 5’-GCAAGAACCTGATGGACATGTTCAG-3’ and 5’-GCAATCCCCAGAAATGCCAGATTAC-3’

### Preparation of isolated pancreatic islets from mice

Pancreatic islets were isolated as previously described (Doliba et al., 2017; Li et al., 2009). For each experiment, 1 to 4 adult mice (8 - 12 weeks of age) were euthanized by exposing them to an overdose of isoflurane through inhalation and a subsequent cervical dislocation.

Following a perfusion through the common bile duct with a Hank’s balanced salt solution (HBSS; GIBCO, 14175) containing collagenase XI (1 - 2 mg/mL, Sigma, C7657) and 1 mM CaCl_2_, the pancreas was dissected and digested at 37 °C for 5 or 15 minutes. After the digestion, the collagenase in the pancreas homogenate was removed through several washes with HBSS containing 0.3 % (w/v) bovine serum albumin, and dissociated islets were then hand-picked into a recovery media RPMI1640 (GIBCO, 21870) with the following supplements: 10 % fetal bovine serum, 2 mM GlutaMAX, 1 mM sodium pyruvate, penicillin-streptomycin, and 10 mM HEPES titrated to pH 7.4 with NaOH. For the perifusion assay, the digested pancreases samples were subject to a Ficoll gradient purification, and the islets in the enriched fraction were washed with the recovery solution before handpicking islets. During this process, the samples from four mice were pooled to ensure a sufficient number of islets were used to perform assays under compared conditions with the same population in each independent round of experiment. The handpicked islets were placed in 5 % CO_2_ incubator at 37 °C to rest overnight prior to the static incubation assay or two days prior to the perifusion assay.

### Assays of secreted insulin and total insulin content of mouse islets

For insulin secretion in each independent static incubation assays, pancreatic islets from 1 mouse (8 - 12 weeks of age) were used as follows. Isolated islets were washed by six sequential transfers to a modified Krebs-Ringer buffer containing 114 mM NaCl, 24 mM NaHCO_3_, 5 mM KCl, 2.2 mM CaCl_2_, 1 mM MgCl_2_, 1 mM NaH_2_PO_4_, 10 mM HEPES titrated to pH 7.4 with NaOH, 0.3 % w/v BSA, and 3 mM glucose. Washed islets were equilibrated in the final wash step for one hour at 5 % CO_2_ and 37 °C. Using a previous study as reference (Remedi et al., 2009) and on the basis of the amount of insulin secretion and the sensitivity of insulin assay, we chose to use 5 – 10 islets in each assay. After one hour of equilibration, each group of 5 to 10 randomly selected islets was placed into a well of a micro-well plate containing the modified Krebs-Ringer buffer, added with additional test reagents: glucose and SpTx1 as indicated in Figure 2. Diazoxide (Sigma, D9035) was added to the modified Krebs-Ringer buffer 20 minutes prior to the addition of other test reagents as indicated in Figure 11. Following 1 hour incubation under the conditions of 5 % CO_2_ and 37 °C, the supernatant containing released insulin was recovered and the islets were placed in acidified ethanol to extract total insulin content as previously described (Leiter, 2009). Insulin in the samples was assayed using a colorimetric ELISA kit (ALPCO, 80-INSRTU-E10). The variations in data values reflected both biological variability and technical errors. No experiments were excluded.

For each perifusion experiment (Doliba et al., 2017), pancreatic islets from 4 mice (8 - 12 weeks of age) were used in the following manner. About 180 handpicked islets were placed onto a polycarbonate membrane (Nuclepore, Whatman) inside a polypropylene perifusion chamber (Swinnex, Millipore) and the oxygenated perifusion solution (modified Krebs-Ringer buffer as above) added with additional test reagents: glucose, glibenclamide (Sigma, G0639) and/or SpTx1 as indicated in Figures 9 and 10. All experiments were performed at 37 °C with an HPLC-controlled flow rate of 1 mL/min; consecutive fractions of 1 mL were collected. At the end of each experiment, the islets were recovered and placed in acidified ethanol to extract total insulin content as described above. Insulin in the samples was assayed using an insulin-specific radioimmunoassay kit (RI-13K, Millipore). For each condition, the study was performed 4 to 6 times. The variations in data values reflected both biological variability and technical errors. No experiments were excluded.

### Administration of tamoxifen and induction of diabetes in NDM-model mice

In ^Rosa26^mKir6.2^NDM^ and ^Rosa26^mKir6.2^NDM-V108E^ mice, the expression of mutant Kir6.2 activity occurred in the pancreas upon the excision of the Neo/WSS cassette by the tamoxifen-dependent Cre recombinase (Remedi et al., 2009). Exposure to tamoxifen permitted the translocation of Cre recombinase from the cytoplasm to the nucleus in order to perform the excision between *loxP* sites within the targeted *Rosa26* locus. To induce the expression of mutant Kir6.2 activity and consequently the diabetic phenotype in these mutant mice (8 - 12 weeks old), corn oil containing tamoxifen (75 mg/kg body weight; prepared as described below) was injected intraperitoneally (IP) for five consecutive days. Injection sites were sealed with a tissue adhesive (3M Vetbond) to prevent leakage of the injected liquid.

To make a liquid stock of tamoxifen (20 mg/mL) for injection, tamoxifen solids (Sigma, T5648) were added into corn oil (Sigma, C8267) in a polypropylene tube wrapped with aluminum foil to protect tamoxifen from light. The stock was placed on a nutator to dissolve overnight at room temperature. The dissolved tamoxifen stock was filtered through a sterilized Steriflip unit (Millipore) then aseptically aliquoted, stored at 4 °C, and used within 5 days.

### Mouse blood glucose monitoring and plasma insulin assay

We evaluated the blood glucose levels of tested mice (8 - 12 weeks of age) using a digital glucometer (Clarity Diagnostics, BG1000). Induced-diabetic mice were tested daily starting from the day after the final administration of tamoxifen until their blood glucose levels became above 600 mg/dL. Following overnight (16 hr) fasting, blood glucose monitoring and plasma collection were then performed on wild-type, transgenic, and diabetic mice at designated time points over a 2-hour observation period in the next morning. Glibenclamide (40 mg/kg body weight) or its vehicle DMSO was administered through a single intraperitoneal injection whereas SpTx1 (1 mg/kg body weight) or its vehicle normal saline solution was intravenously administrated by a lateral tail vein or retro-orbital sinus injection. At each given time point, the glucose level of a blood drop obtained from the cut tail tip was measured twice using a glucometer. For any case where the level was above 600 mg/dL, the blood sample was diluted by an equal volume of the phosphate-buffer solution titrated to pH 7.3 before the glucose level was re-measured for two times. Additionally, 4-5 drops (∼12 μL) of tail vein blood were collected at individual specified time points into a tube containing K_2_EDTA (1.5 mg/mL blood) and centrifuged at 2000x g for 5 min at 4 °C. 5 μL of the resulting plasma was frozen in liquid nitrogen, stored at −20 °C, and assayed using a colorimetric insulin-detecting ELISA kit (CrystalChem, 90080) within a week. Mice were grouped according to genotypes. Using a previous study as reference (Remedi et al., 2009), we determined the number of mice for individual groups. The variations in data values reflected both biological variability and technical errors. No experiments were excluded.

### Statistics

Unless otherwise specified, all data are reported as mean (± SEM). Statistical analyses were performed using software Origin 8.0 (OriginLab) and Igor Pro 8 and 9 (Wavemetrics URL link: https://www.wavemetrics.com). *P* values were calculated using 2-tailed *t*-test (Origin 8.0) and presented in the relevant figure legends purely as descriptive parameters of data.

### Software used to generate figures

The data graphs and statistical analyses in Figures 1-12 and S2 as well as data values in Table 1 were created using Origin 8.0 (OriginLab; URL link: https://www.originlab.com). Islet cell electrophysiological results were analyzed using Igor Pro 8 and 9. Snippets of the DNA sequencing chromatograms in Figure S1 were generated by screen capture, the textual contents of the figure legends were drafted with Word (Microsoft Office Suite version 365; https://www.micro soft.com/en-us/store/apps/windows). All figure panels and legends were made by importing and resizing these vector graphics, image files and texts using Adobe Illustrator (version CS4; https://www.adobe.com).

## Supplemental Figure Legends

**Figure S1.**
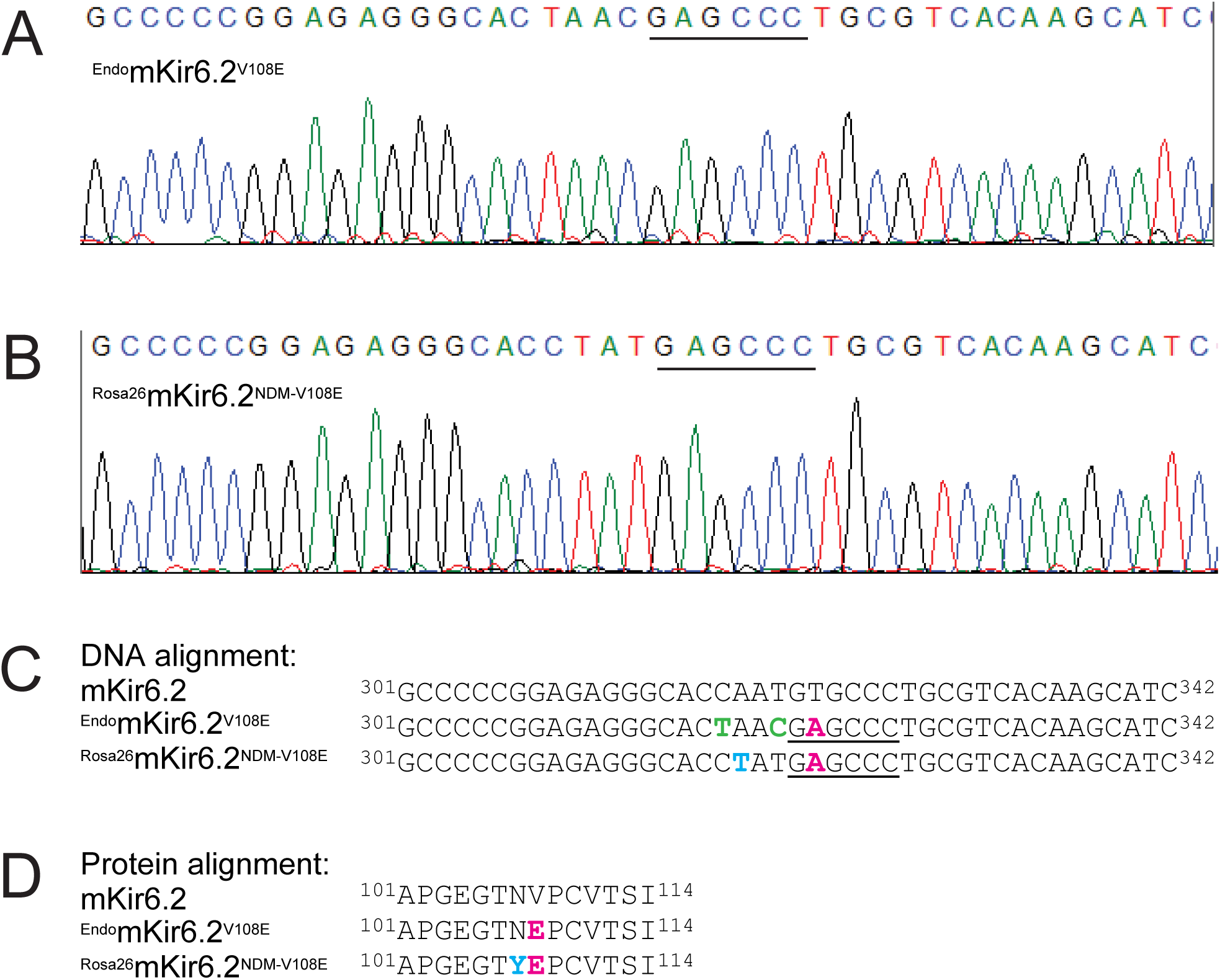
The DNA sequencing results of ^Endo^mKir6.2^V108E^ and ^Rosa26^mKir6.2^NDM-V108E^ mice. (**A**, **B**) Chromatograms showing the sequencing results of a 42-nucleotide segment within the PCR-amplified products of mKir6.2 in the endogenous (A) or the *Rosa26* (B) locus; the engineered-in restriction enzyme (BanII) cleave site in each is indicated by the black line (Methods). (**C**) Partial DNA sequence of wild-type Kir6.2 aligned with those, shown in **A** and **B**, from the mutant Kir.6.2 in the endogenous or the *Rosa26* locus. Intended non-synonymous or synonymous mutations are colored magenta or lime whereas an unintended non-synonymous mutation is colored blue; the BanII cleave site is underlined. The Kir6.2 gene at the endogenous locus contained three intended mutations in the DNA that include: a non-synonymous substitution of T323A (magenta) in the targeted DNA site that causes the valine-to-glutamate mutation at position 108 in the protein to confer higher SpTx1-sensitivity on the mouse Kir6.2 channel protein; and two synonymous substitutions of C318T and T321C (lime) in the targeted DNA site to prevent re-editing by Cas9 while preserving the original amino acids threonine 106 and asparagine 107 in the Kir6.2 protein. However, the Kir6.2 transgene at the *Rosa26* locus contained the same intended non-synonymous substitution of T323A (magenta) besides an unintended non-synonymous substitution of A319T (blue) in the DNA that leads to an additional mutation of N107Y in the Kir6.2 protein (see Figure S2). (**D**) Alignment of amino acid sequences, corresponding to the DNA sequences (**C**), are aligned among wild-type Kir6.2, mutant Kir.6.2 at the endogenous, and mutant Kir.6.2 *Rosa26* locus, where the intended V108E mutation or unintended N107Y mutation is colored magenta or blue.

**Figure S2.**
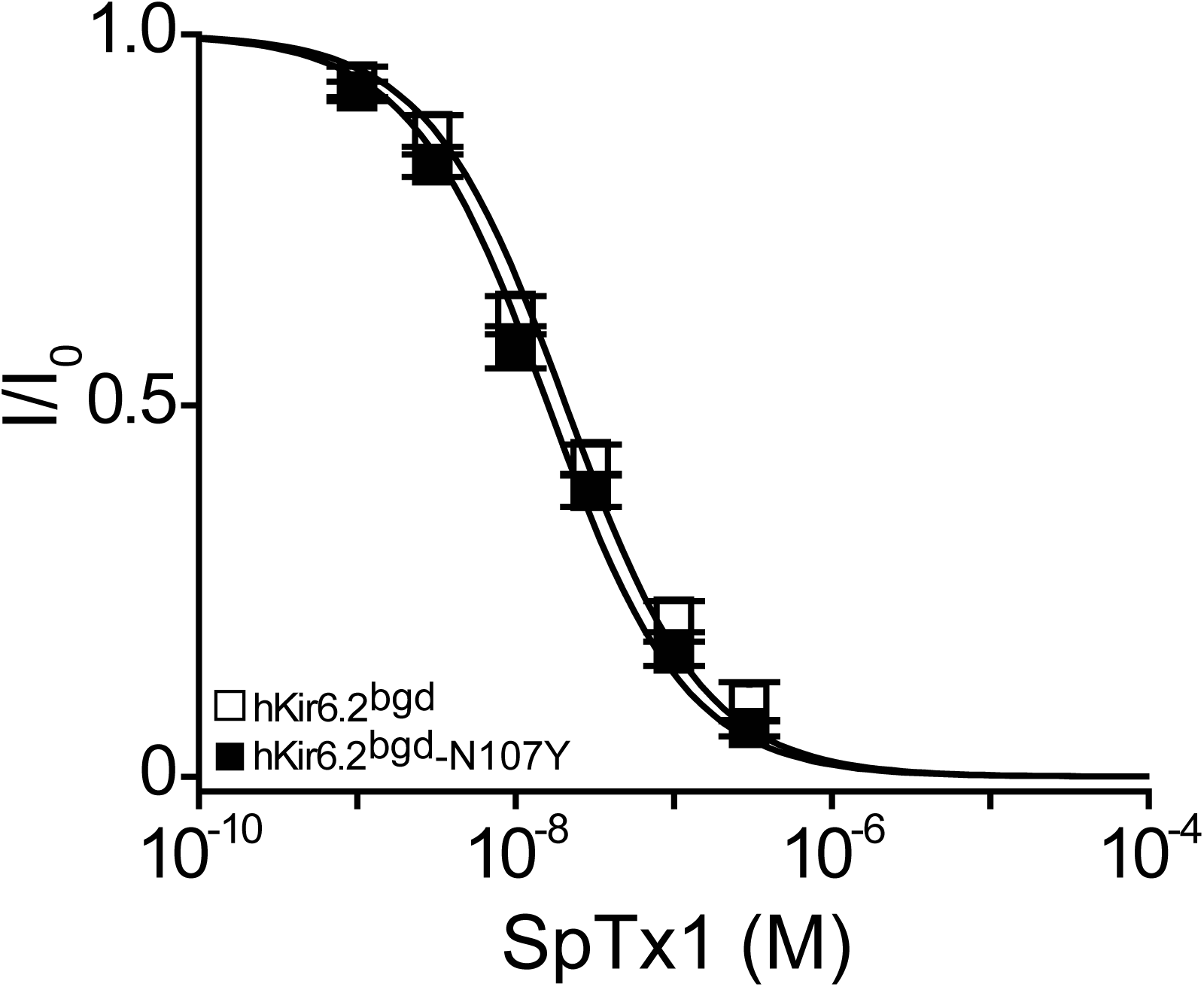
The N107Y mutation does not reduce the SpTx1 sensitivity of hKir6.2^bgd^. Fractions of remaining channel currents (I/I_o_) of hKir6.2^bgd^ without (open squares) or with (filled squares) the N107Y mutation plotted against the concentration of SpTx1. The curves superimposed onto the data correspond to the fits of an equation for a bimolecular reaction. The fitted K_d_ values are 2.08 (± 0.18) × 10^-8^ M for hKir6.2^bgd^, and 1.61 (± 0.14) × 10^-8^ M for hKir6.2^bgd^ containing N107Y, where data are plotted as mean ± SEM (*n* = 3).

